# Single-cell atlas of pig-to-monkey kidney xenotransplantation reveals macrophage chimerism and an IFN-ε orchestrated graft protective immune niche

**DOI:** 10.64898/2026.03.21.713416

**Authors:** Hanfeng Wang, Jianwen Chen, Yuan Chang, Weimin Ci, Xiumeng Hua, Fei Yu, Shujun Yang, Xu Zhang, Jiangping Song, Yang Fan

**Affiliations:** Department of Urology, Chinese PLA General Hospital, Beijing, China; Department of Nephrology, Chinese PLA General Hospital, State Key Laboratory of Kidney Diseases, National Clinical Research Center for Kidney Diseases, Beijing, China; The Cardiomyopathy Research Group at Fuwai Hospital, State Key Laboratory of Cardiovascular Disease, National Center for Cardiovascular Diseases, Beijing, China; Beijing Key Laboratory of Preclinical Research and Evaluation for Cardiovascular Implant Materials, Animal Experimental Centre, Fuwai Hospital, Chinese Academy of Medical Sciences and Peking Union Medical College, Beijing, China; Department of Cardiac Surgery, Fuwai Hospital, National Center for Cardiovascular Diseases, Chinese Academy of Medical Sciences and Peking Union Medical College, Beijing, China

## Abstract

The clinical translation of xenotransplantation is constrained by an incomplete understanding of the cellular circuitry governing graft rejection versus adaptation under current immunosuppressive regimens. Here, we present a high-resolution single-cell atlas of pig-to-monkey kidney xenotransplantation in a preclinical model, capturing the early immune dynamics preceding graft loss. Our analysis reveals a landscape dominated by innate immunity, characterized by species-specific distribution of macrophages. We identify recipient-derived macrophage subsets (ACKR1^+^, FN1^+^, MARCO^+^, IDO1^+^) enriched for multiple immune checkpoint molecules, alongside donor-resident subpopulations (APOE^+^, CCL2^+^, IL-1A^+^) exhibiting pro-inflammatory pathogenic signatures, and leverage their transcriptional profiles to computationally predict candidate therapies (belatacept, abatacept, pexidartinib). Further analysis demonstrates that despite divergent differentiation trajectories, both recipient-derived and donor-resident macrophages converge toward hybrid M1/M2 states at their terminal stages. Notably, we uncover a graft-protective circuit orchestrated by epithelial-derived interferon-epsilon (IFN-ε), which specifically engages multiple subsets including IDO1^+^ macrophages to establish a localized immune-tolerant niche. This study establishes the xenograft epithelium as an active participant in immune modulation via the IFN-ε axis and reveals macrophage functional chimerism as a key actionable feature of cross-species immune responses.

## Introduction

Organ transplantation is the definitive therapy for end-stage organ failure, yet the global shortage of donor organs remains a paralyzing barrier, depriving millions of life-saving treatment annually^1^. Xenotransplantation using genetically engineered porcine organs offers a transformative solution, leveraging pigs’ anatomical and physiological similarity to humans, rapid maturation, and scalable breeding^2–4^. While pivotal advances such as α-galactosyltransferase (GGTA1) knockout have mitigated hyperacute rejection, long-term graft survival remains elusive due to unresolved immunological hurdles: acute antibody-mediated (AbMR) and T cell-mediated rejection (TCMR), immune memory, and cross-species immune crosstalk^5–7^. These challenges demand rigorous preclinical models to decipher mechanisms and validate interventions before clinical translation.

Recent brain-dead decedent (BDD) studies, including 61-day extensions^8–11^, have yielded pivotal initial insights into pig-to-human xenotransplantation. They demonstrate that genetically engineered porcine kidneys sustain short-to-mid-term physiological function in anephric humans, supporting urine production, creatinine clearance, and electrolyte balance. BDD models have uncovered early AbMR hallmarks^12,13^, and validated complement inhibition/plasma exchange for rejection reversal^8^, while single-cell with spatial transcriptomics have identified human immune cell infiltration and porcine tissue repair programs^14^. However, inherent limitations restrict their utility: brain death-induced systemic inflammation, cytokine storms, and hypothalamic-pituitary dysfunction confound immune response interpretation^15,16^; even extended follow-up fails to capture long-term endpoints (chronic rejection, immune memory, late-onset antibodies) critical for clinical success; longitudinal adaptive immune monitoring and novel immunosuppressant validation are unfeasible; and compromised renal-immune physiology limits modeling of living patient scenarios^11,15,17^.

By contrast, the pig-to-non-human primate (NHP) model potentially addresses these gaps through irreplaceable strengths: intact physiological immune systems eliminate brain death-related confounders, enabling precise characterization of xenoreactive responses^18–20^; long-term follow-up permits assessment of chronic rejection and immune memory^6^; iterative testing of targeted therapies validates interventions for clinical translation^21^; and evolutionary proximity to humans ensures conserved immune pathways with superior predictive value^15,22^. These attributes have established NHPs as the primary model for preclinical xenotransplantation.

Despite significant progress, the precise single-cell dynamics governing pig-to-NHP xenograft rejection remain poorly defined. Here, we apply single-cell transcriptomics to pig-to-monkey renal xenografts to delineate intra-graft immune remodeling, xenoreactive cell subsets, and species-specific molecular interactions, identifying unrecognized rejection drivers and providing a high-resolution roadmap for optimizing graft design and targeted immunomodulation. The data resources from this study are expected help deepen understanding of xenotransplantation immune rejection mechanisms and provide modest support for clinical translation of genetically engineered porcine organs.

## Results

### Single-cell profiling reveals innate immune dominance in xenotransplant kidneys under immunosuppression

To delineate cell-type-specific responses in xenografts under immunosuppression targeting adaptive immunity, we employed a kidney xenotransplantation model using two α-GAL-knockout (GGTA1 monoallelic KO) Bama miniature pigs as donors (male, 67-68 days old) and two rhesus macaques as recipients (male, 10-11 years old). Recipients received a clinically mimetic dual-phase immunosuppression regimen: induction with basiliximab, methylprednisolone, and rituximab, followed by maintenance therapy with mycophenolate mofetil, tapered corticosteroids, and cyclosporine A. Xenograft kidneys were collected at terminal rejection (post-transplant day 7 and day 11; n=2) along with untransplanted α-GAL-KO pig kidneys as controls.**(Fig. 1A, Supplementary Table 1)**. Single-cell sequencing samples from all kidneys were obtained following standardized criteria for anatomical region, depth, and tissue volume.

**Figure 1.**
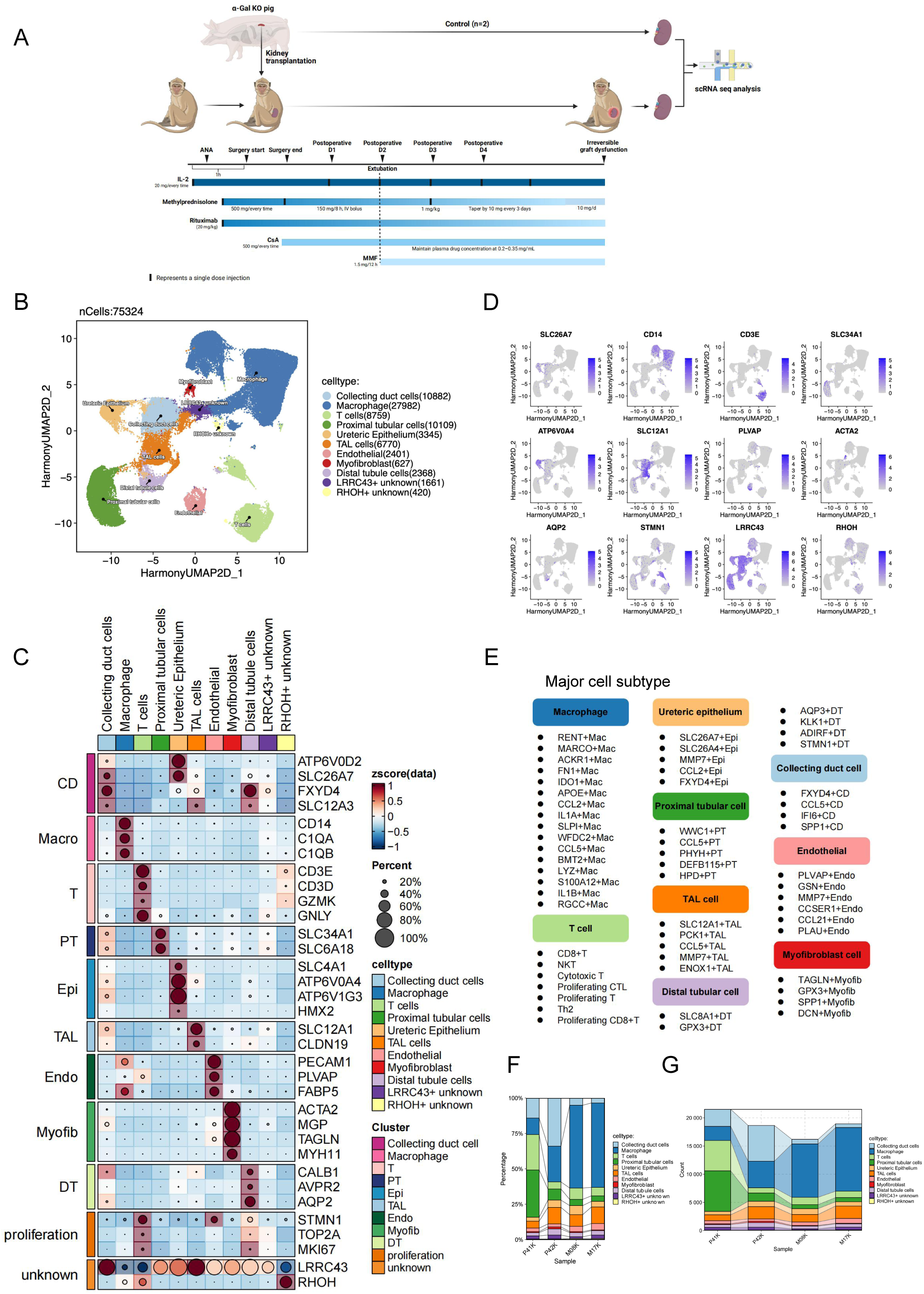
Cellular landscape of pig-to-monkey kidney xenografts. (A) Schematic overview of the study design and workflow. (B) UMAP visualization of major clusters across all samples. (C) Bubble heatmap showing expression levels of selected signature genes across all major cell populations. Dot size indicates fraction of expressing cells, colored based on normalized expression levels. (D) UMAP visualization of marker genes used for identifying cell types in single-cell RNA sequencing datasets. (E) The subsets of each major cell populations with labeling signature genes. (F) Relative proportion of all major cell populations in individual samples. (G) Cell counts of all major cell populations in individual samples.

After rigorous quality control **(Extended Data Fig. 1A, 1B)**, 75,324 high-quality cells were analyzed. Unsupervised clustering identified nine transcriptionally distinct major populations, including renal epithelial (collecting duct [CD], proximal tubule [PT], thick ascending limb [TAL], distal tubule [DT], ureteric epithelium [Epi]), stromal (endothelial [Endo], myofibroblasts [Myofib]), and immune cells, alongside two unclassified clusters defined by LRRC43 and RHOH overexpression **(Fig. 1B-1D, Supplementary Table 2)**. Subclustering resolved 58 functionally specialized subsets with unique spatial distributions **(Fig. 1E, Supplementary Table 3-7)**. Strikingly, under given immunosuppressive regime, we observed pronounced enrichment of general macrophages **(Fig. 1F, 1G, Extended Data Fig. 2A-2E)** in xenotransplant kidneys, indicating aberrant innate immune activation.

### Xenografts exhibit severe rejection injury

Compared to the control kidneys **(Fig. 2A, sections a and c)**, both xenotransplanted kidneys exhibited mild-to-severe glomerular **(Fig. 2A, sections b1 and d1)** and tubular injury **(Fig. 2A, sections b3 and d3)**. Xenograft 2 demonstrated the most significant injury, with conspicuous focal necrosis **(Fig. 2A, sections d1 and d3)**. Concurrently, microvascular endothelial injury with marked interstitial hemorrhage **(Fig. 2A, sections b2 and d2)** was evident in both xenografts, although microthrombosis were not observed. Furthermore, both xenografts developed prominent microvascular inflammation (MVI), characterized by glomerulitis and peritubular capillaritis, that was most severe in the graft from Recipient 1 (day 7). The pathologic diagnosis of antibody-mediated rejection and the complement system activation was molecularly confirmed by strong, diffuse glomerular deposition of IgM, IgG and complement C4d **(Fig. 2B)**. In contrast, endothelial markers CD31 and von Willebrand factor (vWF) showed only weak staining, with no significant difference from controls. Consistent with the observed innate immune dominance, CD68^+^macrophages exhibited aberrant chemotactic enrichment and displayed a diffuse distribution pattern throughout the xenograft interstitium **(Fig. 2C)**. This was accompanied by a sharp decline in CD3^+^ lymphocytes and a marked increase in CD15^+^innate immune cells **(Extended Data Fig. 3A)**.

**Figure 2.**
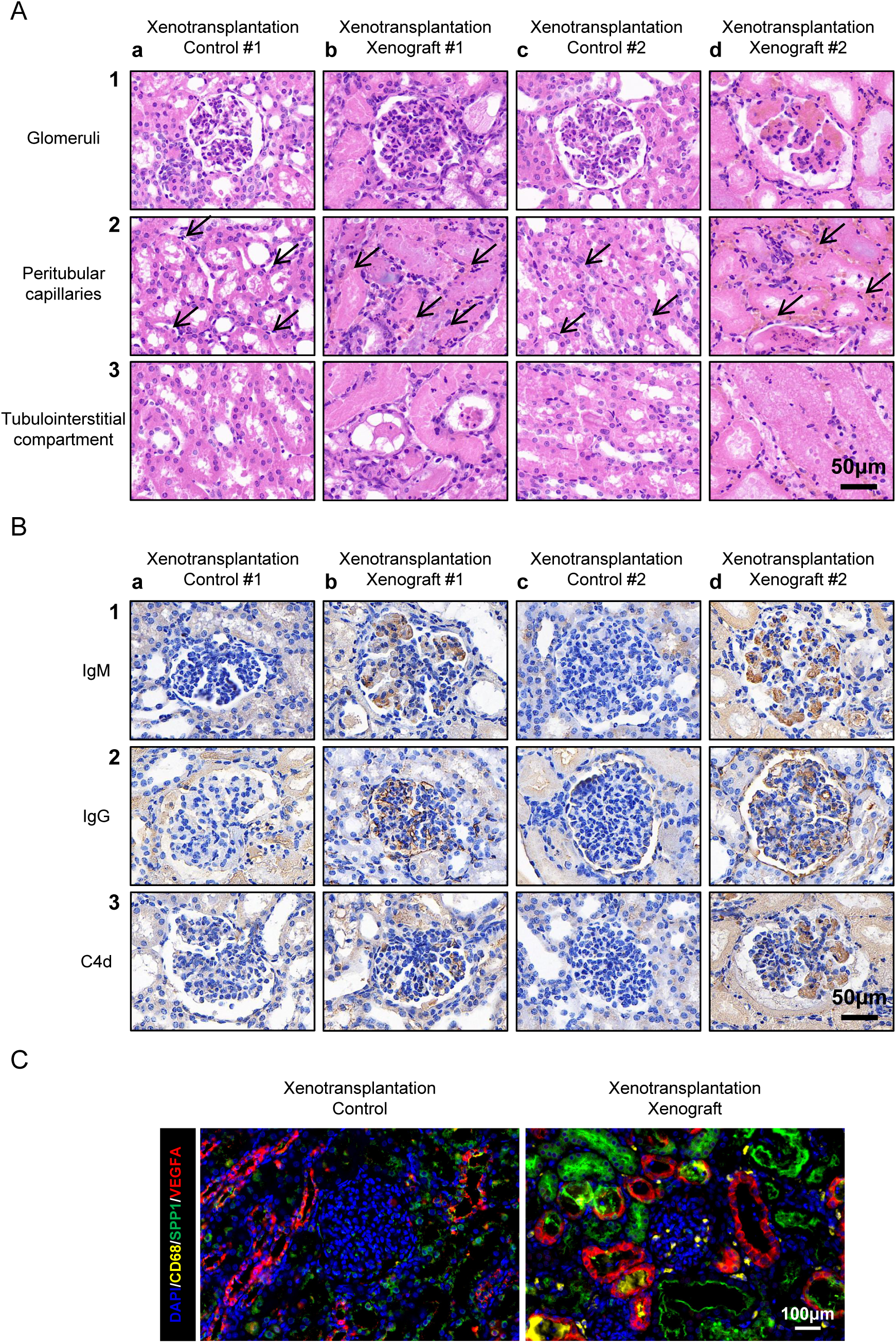
Histopathological features in xenograft. (A) Haematoxylin and eosin-based evaluation for xenotransplantation control #1 (sections a1-a3) and #2 (sections c1-c3), xenograft #1 (sections b1-b3) and #2 (d1-d3). Fields representative of glomeruli (row 1), peritubular capillaries (row 2), and tubulointerstitial compartments (row 3) are shown. Black arrows show non-infiltrated peritubular capillaries (sections a2 and c2), and infiltrated peritubular capillaries (section b2 and d2). (B) Immunohistochemistry-based evaluation for xenotransplantation control #1 (sections a1-a3) and #2 (sections c1-c3), xenograft #1 (sections b1-b3) and #2 (d1-d3). Fields representative of IgM (row 1), IgG (row 2) and C4d (row 3) stainings are shown. (C) Immunofluorescence staining of infiltrated macrophages with signature proteins. Scale bars, 100 µm.

### Myeloid remodeling in xenografts reveals divergent donor and recipient dynamics

Beyond 16 transcriptionally distinct macrophage sub-clusters, unsupervised clustering revealed two canonical myeloid lineages, ISG15^+^ neutrophils and FLT3^+^ dendritic cells (DCs), based on established markers **(Fig. 3A, 3B)**. Additionally, two unclassified myeloid sub-clusters emerged: a proliferative cluster (marked by STMN1, MKI67 and UBE2C), and a pro-inflammatory cluster (defined by PLAVP, ADGRG1, and PTGS1) **(Fig. 3A, 3B).** To dissect the origin of myeloid cells in the grafts, scRNA-seq reads were independently aligned to porcine (donor) and monkey (recipient) genomes. Each dataset was analyzed independently to minimize technical variability and ensure robust characterization of myeloid subsets. Four macrophage subsets (MARCO^+^, ACKR1^+^, FN1^+^, IDO1^+^) were exclusively recipient-derived, and APOE^+^, CCL2^+^, IL1A^+^, SLPI^+^, WFDC2^+^ and CCL5^+^ remained all of donor origin. **(Fig. 3C, Extended Data Fig. 4A, 4B)**.

**Figure 3.**
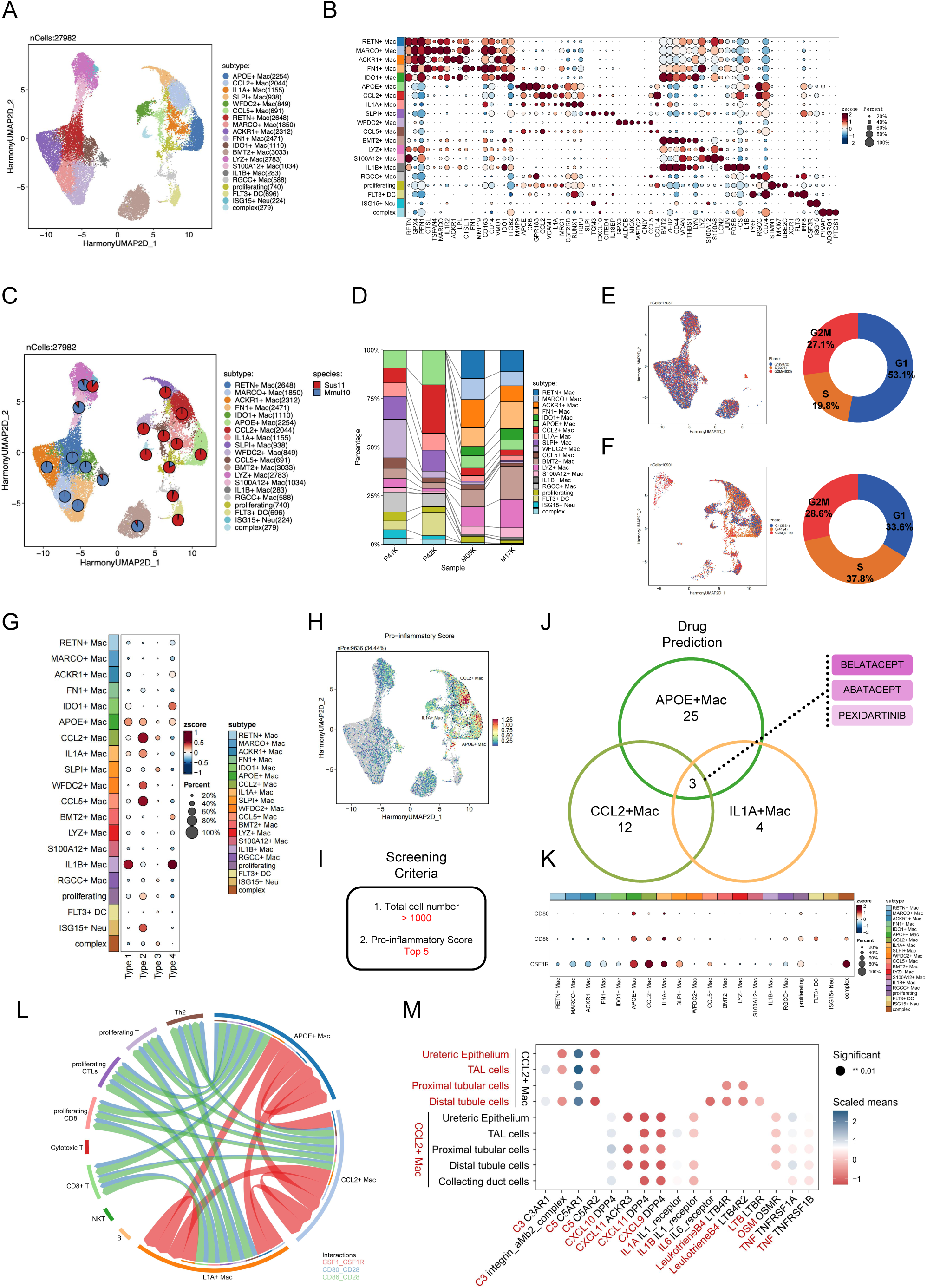
Myeloid landscape identifies targetable pro-inflammatory subsets. (A) UMAP visualization of myeloid cell lineages across all samples. (B) Bubble heatmap showing expression levels of selected signature genes across myeloid cell lineages. Dot size indicates fraction of expressing cells, colored based on normalized expression levels. (C) UMAP plot depicting the distribution of cells from different species within a specific subclusters. (D) Relative proportion of myeloid cell lineages in individual samples. (E-F) UMAP plots showing the cell cycle assignment of myeloid cell lineages derived from recipient and tissue-resident respectively. (G) Type 1-4 pro-inflammatory scores (type 1: TNF, IL-1A, IL-1B, IL-6; type 2: CXCL8, CCL2, CX3CL1, CCL5, CCL11; type 3: IFNB, CXCL9, CXCL10, CXCL11; type 4: IL-1B, IL-18) for the identified macrophage substes. (H) Combined type 1-4 pro-inflammatory expression score per cell overlayed on the UMAP projection. The positions of CCL2^+^, APOE^+^ and IL1A^+^ macrophage subsets are highlighted manually. (I) Screening Criteria of macrophage subsets for following durg prediction. (J) Quantification of predicted drug-macrophage subpopulation interactions. (K) Bubble heatmap showing expression levels of CD80, CD86 and CSF1R across myeloid cell lineages. Dot size indicates fraction of expressing cells, colored based on normalized expression levels. (L) Circo-plot showing the intercellular interactions between lymphoid cell lineages and CCL2^+^, APOE^+^ and IL1A^+^ macrophage subsets. The strings represented interactions determined based on the expression of a ligand by one cell cluster and the expression of a corresponding receptor by another cell cluster. The string thickness reflects the number of different interaction pairs, colored according to cell cluster. (M) Dot plots showing L-R interactions between CCL2^+^ macrophages and TECs. P values were calculated using an empirical shuffling test.

Critically, we observed a pronounced decline in tissue-resident macrophage populations (including CCL2^+^, SLP1^+^, WFDC2^+^) following transplantation. This atrophy extended to resident-derived ISG15^+^ neutrophils and FLT3^+^ DCs (**Fig. 3D, Extended Data Fig. 4A, 4B**), indicating a broad functional impairment of the porcine myeloid compartment under xenogeneic stress. Supporting this interpretation of functional compromise, cell-cycle profiling uncovered divergent proliferative dynamics: recipient-derived macrophages skewed towards G1 phase (>50%) while donor-derived macrophages exhibited a disproportionately elevated S-phase proportion relative to G1/G2M phases, suggesting impaired proliferative capacity (**Fig. 3E, 3F**).

### Drugable pro-inflammatory macrophage subsets identified

Given the pronounced enrichment of macrophages and their stark numerical disparity between donor and recipient origins, we hypothesized that specific pro-inflammatory subsets mediate xenograft rejection. To test this, we characterized cytokine expression profiles across myeloid clusters.

Single-cell analysis identified IL-1B⁺ and APOE⁺ macrophages as core producers of Type 1 cytokines (TNF, IL-1A/B, IL-6). Type 2 inflammatory mediators (CXCL8, CCL2, CX3CL1, CCL5, CCL11) were broadly expressed across multiple subsets, including APOE⁺, CCL2⁺, IL-1A⁺, CCL5⁺, and WFDC2⁺ populations. In contrast, Type 3 mediators (IFNB, CXCL9/10/11) were scarcely detectable, while Type 4 cytokines (IL-1B, IL-18) were predominantly enriched in IDO1⁺ and IL-1B⁺ macrophages **(Fig. 3G)**.

To prioritize the most pathologically relevant pro-inflammatory subsets, we applied a dual filter: clusters containing >1,000 cells and ranked among the top 5 by pro-inflammatory score. This identified three donnor-derived macrophage subsets, APOE⁺, CCL2⁺, and IL-1A⁺, which exhibited the strongest and broadest pro-inflammatory signatures, establishing them as primary therapeutic targets **(Fig. 3H-J, Supplementary Table 8)**. A subsequent transcriptome-guided digital pharmacology screen of FDA-approved compounds identified belatacept, abatacept, and pexidartinib as promising candidates that specifically engage all three pathogenic macrophage subsets **(Fig. 3J)**, highlighting their potential for therapeutic repurposing.

To assess the translational potential of the prioritized drugs, we profiled the expression patterns of their respective targets across all myeloid subsets. The cognate receptors and ligands for belatacept/abatacept (CD80, CD86) and pexidartinib (CSF1R) were preferentially enriched in the APOE⁺, CCL2⁺, and IL-1A⁺ macrophage subsets compared to other myeloid populations **(Fig. 3K)**. Ligand-receptor (L-R) interaction analysis further substantiated these findings, revealing strong inferred crosstalk between CD80/CD86 on these macrophage subsets and CD28 on lymphoid cell lineages, especially T cells, as well as between macrophage CSF1R and its cognate ligands CSF1 expressed by distinct cellular compartments **(Fig. 3L, Extended Data Fig. 4C)**. Beyond these immune interactions, in-depth L-R analysis uncovered that all three pathogenic macrophage subsets engage in extensive, pro-inflammatory signaling networks with tubular epithelial cells (TECs), involving pathways associated with epithelial injury and tissue damage (IL6-IL6R, TNF-TNFRSF1A/B, CXCL9/10/11-DPP4) (**Fig. 3M, Extended Data Fig. 4D**).

### IFNE^+^ TECs orchestrate a graft-protective microenvironment

IFN-ε (IFNE) is a potent immunomodulator ^23^. Three type I interferons (IFNB1, IFNE, IFNK) and one type II interferon (IFNG) were identified in our xenogeneic kidney transplantation model, with striking expression profile differences: all four were expressed in macrophages; IFNG was most abundant in T cells, where IFNK was undetectable. Notably, both IFNG and IFNE were expressed across all 11 cell populations, albeit at variable levels **(Fig. 4A, Extended Data Fig. 5A)**. Intergroup comparisons showed significant upregulation of the IFNG pathway in xenograft T cells, consistent with T cell-mediated immune rejection activation, while the pathway was markedly downregulated in macrophages, CD, PT, Epi, TAL, and Endo cells. IFNE exhibited a unique expression signature: no statistical difference in CD, PT, or Epi cells; downregulated yet detectable expression in TAL cells; and, notably, significant upregulation (rather than decrease) in DT cells **(Fig. 4B)**.

**Figure 4.**
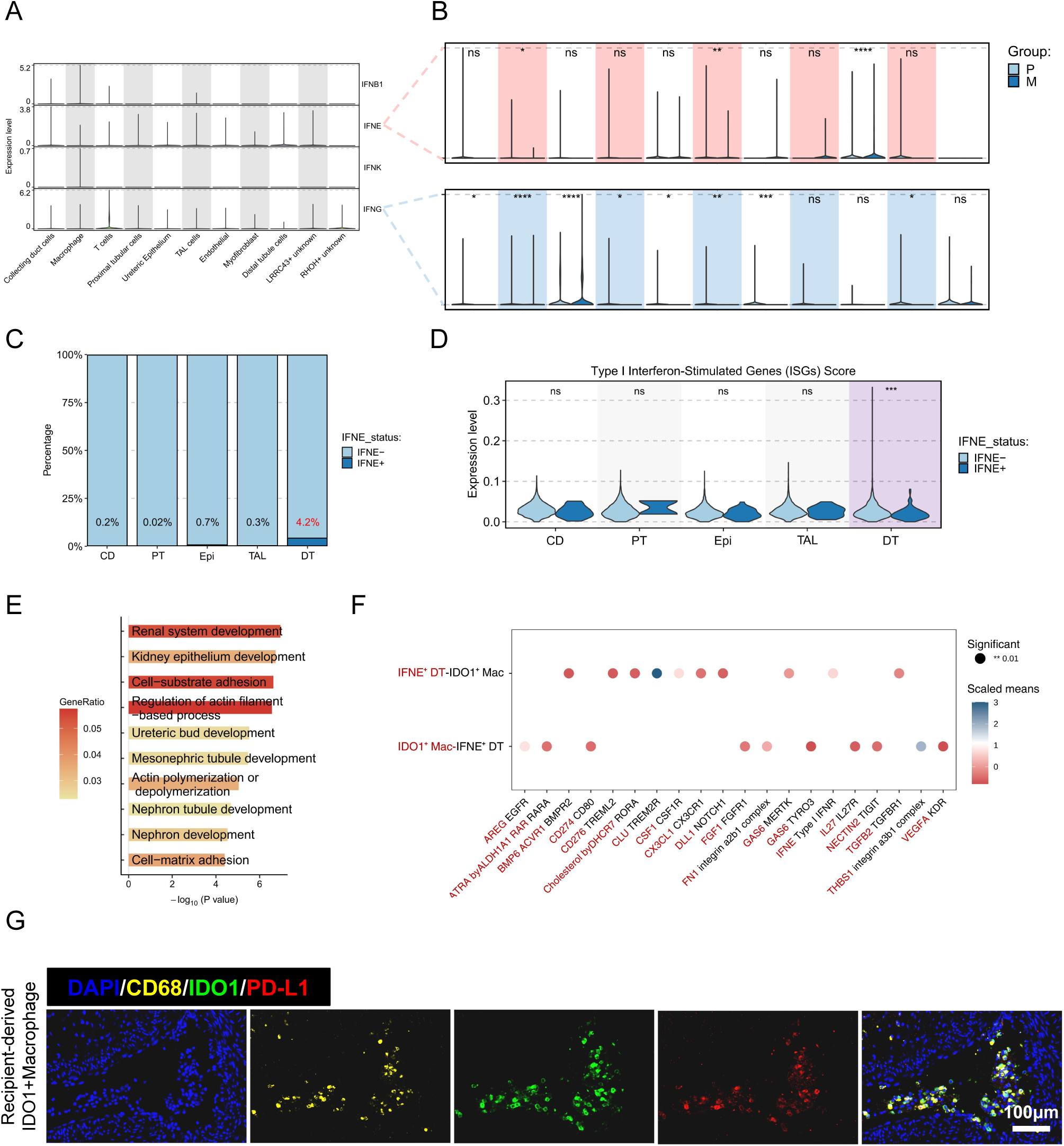
Transcriptional features and intercellular crosstalk of IFNE^+^ tubular epithelial cells (TECs) (A) Violin plots showing interferon expression profiles across major cell clusters. (B) IFNE and IFNG expression across major clusters in control versus xenograft groups. (C) Distribution of IFNE^+^ cells among TEC clusters. (D) Type I ISG module scores in IFNE^+^ versus IFNE^−^ TECs across all clusters. (E) GO enrichment analysis of genes upregulated in IFNE^+^ TECs compared to IFNE^−^TECs. (F) Dot plots showing L-R interactions between IFNE^+^ distal tubule cells and IDO1^+^ macrophage. P values were calculated using an empirical shuffling test. (G) Immunofluorescence staining of IDO1^+^ macrophages with signature proteins. Scale bars, 20 µm.

To dissect the functional role of the IFNE pathway in TECs, we labeled IFNE^+^ cells and found they constituted up to 4.2% of DT cells, with substantially higher type I interferon-stimulated gene (ISG) scores compared to IFNE^−^ cells **(Fig. 4C, 4D, Extended Data Fig. 5B, Supplementary Table 9)**. Differential gene enrichment analysis revealed that upregulated genes in IFNE^+^ cells were enriched in pathways including renal tubule/nephron development, actin cytoskeleton regulation, and cell adhesion, whereas downregulated genes were enriched in mitochondrial energy metabolism, p53-mediated apoptosis, and meiosis-related pathways **(Fig. 4E, Extended Data Fig. 5C)**. Notably, IFNE^+^ DT cells engaged in extensive crosstalk exclusively with themselves, macrophages, and T cells, contributing to local microenvironmental homeostasis and tissue repair **(Extended Data Fig. 5D)**. Among these, IFNE^+^ DT cells specifically regulated recipient-derived IDO1^+^ immunosuppressive macrophages, establishing a transplanted kidney protective microenvironment centered on immune tolerance, underpinned by protective macrophage recruitment, and supported by renal tubule repair and antifibrosis. Recipient-derived IDO1^+^ macrophages actively protected donor IFNE^+^ DT cells through multiple signals, immune checkpoints, anti-inflammatory, pro-repair, and metabolic regulation **(Fig. 4F, 4G)**. This cell-cell crosstalk may represent a key mechanism whereby donor epithelium actively suppresses rejection and extends graft survival in xenotransplantation, underscoring the pivotal role of IFNE^+^ epithelial cells in xenogeneic immune tolerance.

### Developmental reprogramming drives recipient macrophage divergence

In contrast to the classical M1/M2 dichotomy defined by distinct transcriptional programs ^24^, here we identify recipient-derived macrophage subsets in xenografts that exhibit a hybrid polarization state. Notably, RETN^+^ and FN1^+^ clusters showed significant a weak correlation between M1 and M2 scores (r=0.27, p<2.2e-16) **(Fig. 5A, Extended Data Fig. 6A, Supplementary Table 8)**. Functional profiling further supports this plasticity, revealing that ACKR1^+^, RETN^+^, and MARCO^+^ subsets co-express signatures of both phagocytic and pro-angiogenic programs **(Fig. 5B, Extended Data Fig. 6B, Supplementary Table 8)**.

**Figure 5.**
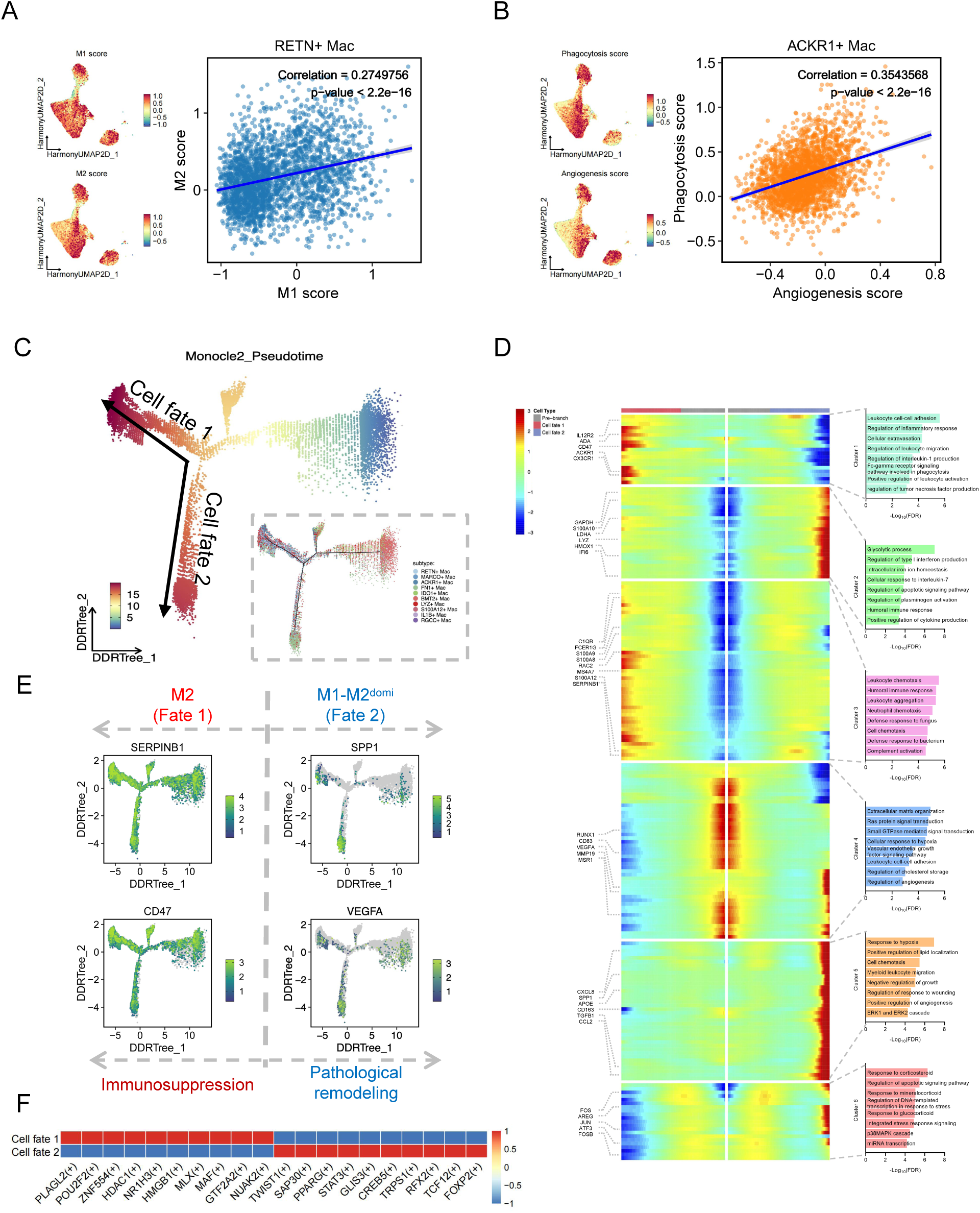
Bifurcated differentiation paths of recipient macrophages. (A-B) UMAP visualization of M1/M2, phagocytosis and angiogenesis signature scores in recipient-derived macrophages, with Pearson correlation coefficients for representative subclusters. (C) Inferred developmental trajectory of recipient-derived macrophages. Arrows in the main plot revealed a dichotomic cell fate division. (D) Heatmap depicting the dynamic expression changes of genes and related pathways of recipient-derived macrophages with different cell fate along pseudotime. (E) Trajectory plots with color gradients indicate dichotomic functional division with the expression level of signature genes (SERPINB1 and CD47 for immunosuppression, SPP1 and VEGFA for tissue remodeling). (F) Heatmap showing transcription factor activity for recipient-derived macrophages with different cell fate.

To explore the putative ontogeny of this hybrid state, we performed pseudotemporal trajectory analysis. This analysis reconstructed a potential differentiation trajectory, which suggested that recipient-derived macrophages could arise from a putative progenitor-like State 1^25^, enriched for small GTPase signaling (SIPA1L1, RIPOR2) and mTOR pathway regulation (BMT2, SIK3), into three terminal branches **(Fig. 5C, Extended Data Fig. 6C, 6D)**: an effector branch (State 2) high in phagocytosis regulators (LYN, LYST) and immune activation markers (SPI1, ITGAL); a metabolic branch (State 3) dominated by oxidative phosphorylation genes (NDUFA1, NDUFB6), aligning with an M2-like, anabolic phenotype^26^; and remodeling branches (States 4-5) featuring ribosomal biogenesis or hypoxia-responsive effectors (VEGFA, MMP14) **(Extended Data Fig. 6E, Supplementary Table 10)**.

Branched expression analysis confirmed the functional divergence of these fates **(Fig. 5D, 5E, Extended Data Fig. 6F).** Fate 1, exhibiting a canonical M2-like profile, established an immune-quiescent niche via CD47/ACKR1-mediated inhibition of phagocytic activity and chemokine recruitment and ADA/SERPINB1-mediated anti-inflammatory cascades, alongside impaired antigen presentation (CD83/MSR1). In contrast, Fate 2 engaged in pathological tissue remodeling via pro-angiogenic and fibrotic programs. Notably, key inflammatory chemokines (CXCL8, CCL2) were sustained across this branch, underpinning a pervasive pro-inflammatory microenvironment that further evidences a hybrid M2-dominant/M1-coupled activation state (M1-M2^domi^), wherein M2-dominant transcriptional programs are coupled with persistent M1-associated inflammatory damage..

Regulatory network analysis identified the core transcription factors governing this fate decision **(Fig. 5F)**. Fate 1 homeostasis was maintained by NR1H3 and HDAC1, whereas Fate 2 pathological remodeling was driven by STAT3 and TWIST1. This bifurcated transcriptional circuitry elucidates the molecular basis for the context-dependent plasticity of recipient-derived macrophages, enabling their adaptation to a rejection microenvironment beyond rigid M1/M2 classifications.

### Metabolic reprogramming shapes resident macrophage hybrid statess

Tissue-resident macrophages critically shape recipient immune responses toward transplanted kidneys ^27,28^. To delineate the functional adaptation of donor-derived tissue-resident macrophages in the xenografts, we first quantified their abundance. Notably, tissue-resident macrophages constituted a major parenchymal population (10,901 cells), rivaling proximal tubule (10,109) and collecting duct (10,882) cells and surpassing total graft-infiltrating T cells (8,759), highlighting their potential to shape the local immune landscape **(Fig. 1B)**.

To investigate their potential adaptation dynamics, we performed pseudotemporal ordering analysis. This inferred a putative developmental trajectory (**Fig. 6A**). The trajectory was rooted in State 4 based on its transcriptional profile, which characterized by humoral immune activation (C1QB, C1QA) and leukocyte differentiation regulation (MAFB, CEBPB). Along this reconstructed path, cells appeared to transition through a metabolic intermediate (State 3) exhibiting a Warburg-like signature (LDHA, PGK1), before branching into two terminal states: State 1, exhibiting enhanced oxidative phosphorylation (NDUFA4, SDHA) aligned with an M2-like phenotype; and State 5, a stress-adaptive state marked by mRNA stabilization (SYNCRIP, HNRNPU) and catabolic regulation genes (NUPR1, VPS35) (**Fig. 6A, Extended Data Fig. 7A-7C, Supplementary Table 11**).

**Figure 6.**
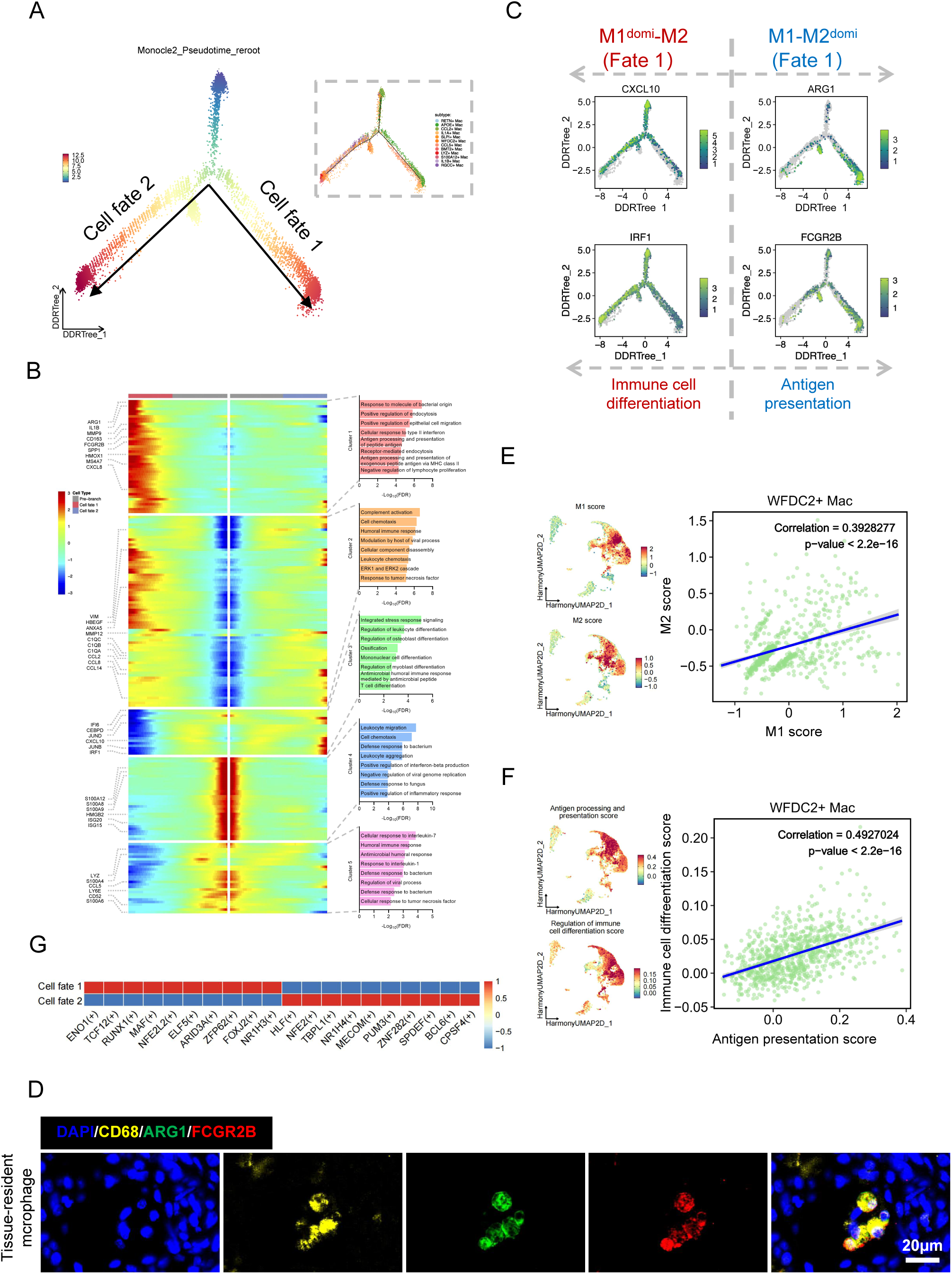
Tissue-resident macrophages acquire hybrid functional states. (A) Inferred developmental trajectory of tissue-resident macrophages. Arrows in the main plot revealed a dichotomic cell fate division. (B) Heatmap depicting the dynamic expression changes of genes and related pathways of tissue-resident macrophages with different cell fate along pseudotime. (C) Trajectory plots with color gradients indicate dichotomic functional division with the expression level of signature genes (CXCL10 and IRF1 for M1^domi^-M2 phenotype of immune cell differentiation, ARG1 and FCGR2B for M1-M2^domi^ phenotype of tissue remodeling). domi=dominant. (D-E) UMAP visualization of M1/M2, antigen processing and immune cell differentiation signature scores in tissue-resident macrophages, with Pearson correlation coefficients for representative subclusters. (F) Heatmap showing transcription factor activity for tissue-resident macrophages with different cell fate. (G) Immunofluorescence staining of tissue-resident macrophages with signature proteins. Scale bars, 20 µm.

Branched heatmap analysis uncovered complex, hybrid polarization patterns within these differentiation paths **(Fig. 6B-6D, Extended Data Fig. 7D)**. Fate 1 (M1^domi^-M2) combined dominant M1-like inflammatory markers (IL1B, CXCL8) with concurrent M2-associated genes (CD163, MRC1). Conversely, Fate 2 (M1-M2^domi^) was characterized by M2-skewed transcriptional programs co-occurring with key M1-skewed signaling mediators (IRF1, CCL2) and leukocyte differentiation priming (CEBPB). This functional duality, wherein cells simultaneously engage in seemingly opposing programs, was further validated by a positive correlation between antigen presentation and immune regulation module scores particular in WFDC2^+^, CCL5^+^, BMT2^+^ subsets **(Fig. 6E, 6F, Extended Data Fig. 7E, 7F, Supplementary Table 8)**.

Regulatory network analysis elucidated distinct transcriptional control mechanisms underlying these hybrid fates **(Fig. 6G)**. Fate 1 (M1^domi^-M2) appeared governed by synergistic activity of NR1H3 and NFE2L2 driving lipid metabolism, coupled with MAF and ELF5 mediating repair programs. Fate 2 (M1-M2^domi^) involved BCL6 and MECOM activating inflammatory responses, alongside NR1H4 and HLF potentially reinforcing a proliferative repair signature.. These divergent circuits provide a molecular basis for the context-dependent plasticity of tissue-resident macrophages, enabling them to transcend classical polarization paradigms and adopt multifaceted functional identities within the xenograft microenvironment.

### Macrophage signaling hubs orchestrate bifurcated immune networks in xenografts

To systematically map intercellular communication networks in xenotransplanted kidneys, we employed L-R interaction analysis. This revealed a marked expansion in both the diversity and intensity of cell-cell signaling networks compared to control kidneys **(Fig. 7A, 7B)**. Macrophages emerged as the predominant immune hubs engaging all major cell types, particularly DT cells, Endo and Myofib clusters **(Fig. 7C-7E)**. The overall network architecture bifurcated into two major axes: one enriched for pro-inflammatory and co-stimulatory interactions, and another dominated by regulatory and inhibitory pathways **(Fig. 7F)** ^29^ (IL1B-IL1R, TNF-TNFRSF1A/B, CXCL10-CXCR3) and T cell co-stimulation signals (CD80/CD86-CD28, ICOSLG-ICOS). Complement pathway interactions, centered on C3, were also prominent. ^30^. Notably, we observed a potential link between complement and interferon signaling via strong interactions between the complement-associated receptor CD93 and IFNGR1 on macrophages and endothelial cells.

**Figure 7.**
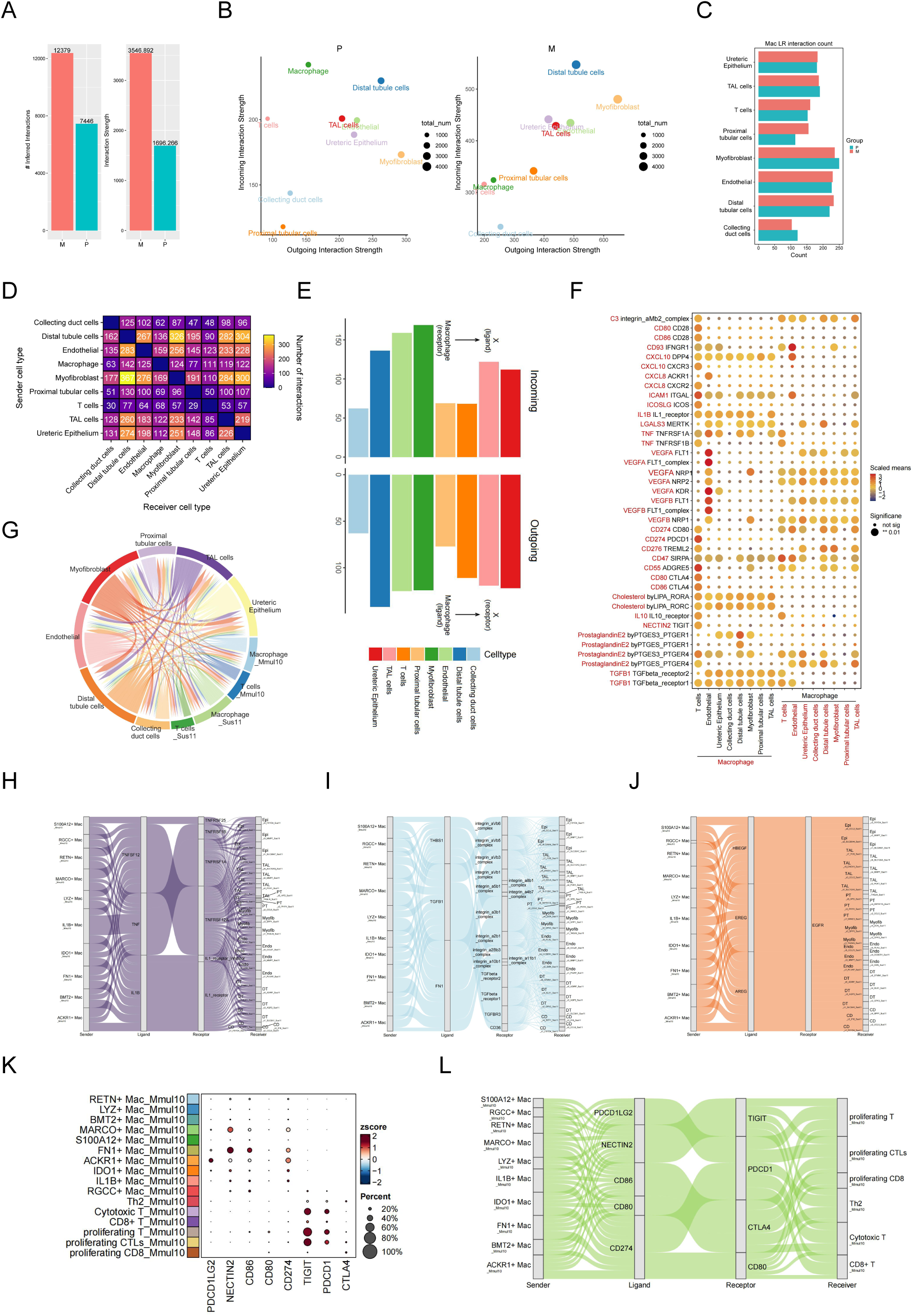
Macrophages exhibited complex cellular interaction network. (A) Total number of inferred ligand-receptor (L-R) interactions and L-R interaction strength for xenotransplantation control and xenograft kidney. (B) Inferred outgoing and incoming L-R interaction strength for individual clusters split by group. (C) Inferred number of macrophage (Mac) L-R interactions with cell clusters stratified by group. Labels refer to clusters. (D) Heatmap showing numbers of potential L-R pairs among indicated cell clusters. (E) Bar plots depicting numbers of putative L-R interactions between Mac and indicated cell clusters. Interaction numbers were calculated based on the expression of receptors and corresponding ligands. Outgoing interactions refer to the sum of ligands from Mac that interact with receptors on the indicated cell cluster. Incoming interactions refer to the opposite. (F) Dot plots showing L-R interactions between macrophage and indicated cell clusters. P values were calculated using an empirical shuffling test. (G) Circo-plot showing the intercellular interactions among indicated cell clusters. The strings represented interactions determined based on the expression of a ligand by one cell cluster and the expression of a corresponding receptor by another cell cluster. The string thickness reflects the number of different interaction pairs, colored according to cell cluster. (H) Sankey diagram showing the pro-inflammatory-specific interactions between recipient-derived macrophages and kidney cells. (I) Sankey diagram showing the fibrogenic-specific interactions between recipient-derived macrophages and kidney cells. (J) Sankey diagram showing the EGFR interactors between recipient-derived macrophages and kidney cells. (K) Dotplot for the ICI genes and subsets highlighted in recipient-derived macrophages and T cells. The size of each dot represents the percentage of cells in the subset expressing the gene. (L) Sankey diagram showing the immunosuppressive-specific interactions between recipient-derived macrophages and T cells for selected ICIs.

Conversely, the regulatory axis featured inhibitory cytokine pathways (TGFB1-TGFbeta_receptor1/2, IL10-IL10R) and immune checkpoint interactions (CD274-PDCD1, CD276-TREML2). This axis also included extensive lipid mediator signaling: prostaglandin E2 (PGE2) signaling via PTGER1/4 receptors polarized macrophages toward M2 phenotypes ^30^, while cholesterol derivatives (LIPA-RORA/RORC) upregulated anti-inflammatory genes (ARG1, IL10) and countered Th17-mediated injury ^31,32^.

Given the varying injury patterns and responses of different TECs clusters under xenogeneic stress, we conducted an in-depth L-R analysis on the subpopulations. As showed in **Fig. 7G, Extended Data Fig. 8A, 8B**, Three recipient-derived macrophage subsets, ACKR1^+^, FN1^+^ and MARCO^+^, showed preferential and specific interactions with injured TEC subpopulations. These interactions implicated distinct mechanistic modules: pro-inflammatory cytokine pairs (IL1B, TNF, TNFSF12, and TNFRSF12A axes) were associated with epithelial damage signatures **(Fig. 7H)**, interactions involving FN1, TGFB1, and THBS1 were linked to fibrogenic pathways **(Fig. 7I)**. Furthermore, a robust network of EGFR signaling interactions (EGFR-EREG/AREG/HBEGF/MIF) from these macrophages to TECs suggested a potential role in driving compensatory epithelial proliferation **(Fig. 7J)**.

These macrophage subpopulations also exhibited pronounced interactions across recipient-derived T cells, particularly proliferating T/CTLs and Th2 clusters **(Extended Data Fig. 8C)**. Among these crosstalks, we noticed that multiple immune checkpoint axes were differentially co-expressed: the ligand NECTIN2 was abundantly expressed on MARCO^+^ macrophages, while FN^+^ macrophages exhibited co-expression of both NECTIN2 and CD86. In contrast, ACKR1^+^ macrophages demonstrated a distinct profile characterized by high expression levels of CD274 and PDCD1LG2. The corresponding receptors TIGIT and PDCD1 were predominantly localized to proliferating T/CTLs and cytotoxic T cell clusters **(Fig. 7K)**. Notably, CD80 was implicated in dual interactions, potentially engaging both the co-stimulatory receptor CD28 and the inhibitory receptor CTLA4 on T cells, as well as acting as a receptor for CD274 **(Fig. 7L)**. This dual targeting may synergistically suppress T cell activation through both PD-1-dependent exhaustion and CTLA-4-mediated co-stimulation blockade. Meanwhile, T cell activation is primarily mediated by immunoglobulin superfamily interactions (ICAM1-ITGAL/integrin_αLβ2 and CD58-CD2) and B7 co-stimulatory signals (CD86/CD80-CD28 and ICOSLG-ICOS) **(Extended Data Fig. 8D)**.

In contrast to the protein-based interactions described above, communication between recipient-infiltrating and donor tissue-resident macrophages was predominantly mediated by lipid signaling molecules **(Extended Data Fig. 8E, 8F)**. Among these, PGE2-PTGER4, Cholesterol/Desmosterol-LXR, and Estradiol-ESR1/GPER1 primarily exert immunoregulatory effects, while DHEA-STS/ESR1 and 5α-DHP-PGR interactions attenuate fibrotic progression. Conversely, LTB4-LTB4R, TXA2-TBXA2R, and LTD4-CYSLTR1/2 predominantly promote pro-inflammatory actions. Collectively, these analyses position macrophages as critical signaling hubs that integrate diverse communication modules to coordinate parenchymal injury, fibrotic remodeling, and adaptive immune responses within the xenograft microenvironment..

### Immunosuppression reshapes lymphocyte composition and stromal interactions

Under the applied immunosuppressive regimen, we collected only 7,057 lymphocytes forming eight sub-clusters, a number far smaller than that of macrophages **(Fig. 8A, 8B)**. Analysis by species origin revealed that recipient-derived lymphocytes outnumbered donor-derived cells by approximately 2.4-fold (**Fig. 8C**). Within this reconstituted immune compartment, donor-derived cells dominated the conventional cytotoxic T lymphocyte (CTL) subset (>90%) and still represented a considerable fraction (>25%) of the proliferating CTL cluster. In contrast, recipient-derived cells constituted the majority in all other lymphoid subsets. Strikingly, donor-derived NKT cells, which were abundant in control kidneys, were nearly undetectable in the xenografts. In addition, CD4^+^ T cell compartment consisted exclusively of Th2-polarized cells. This subset exhibited elevated expression of RGS1, a regulator of GPCR desensitization that may influence Th2 migration ^33^, and of the master transcription factor GATA3, which likely sustains effector functions via the IL-4/IL-5/IL-13 axis^34^. B cells were identified as the smallest immune subpopulation (n=111). While they retained antibody secretory potential, as indicated by expression of JCHAIN, their near-complete numerical depletion compared to controls underscores the efficacy of the anti-CD20-based immunosuppressive strategy (**Fig. 8B-8E, Extended Data Fig. 9A, 9B**).

**Figure 8.**
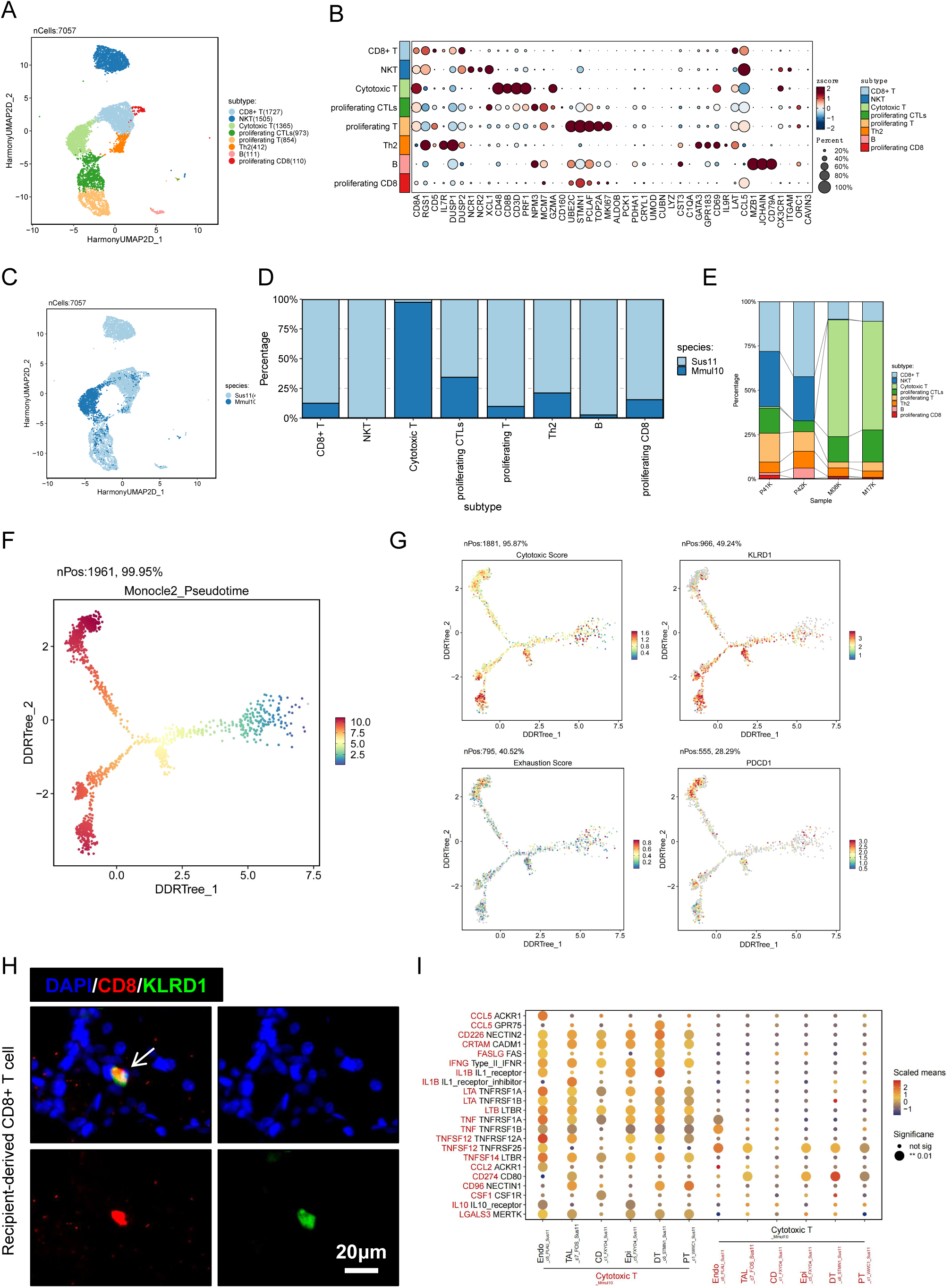
Dynamic variations of T cells at single-cell resolution. (A) UMAP visualization of lymphoid cell lineages across all samples. (B) Bubble heatmap showing expression levels of selected signature genes across lymphoid cell lineages. Dot size indicates fraction of expressing cells, colored based on normalized expression levels. (C) Relative proportion of lymphoid cell lineages in individual samples. (D) Relative proportion of lymphoid cells within xenotransplantation control group (P) and xenotransplantation xenograft group (M). (E) UMAP plot depicting the distribution of lymphoid cell lineages from different species identified in single-cell RNA sequencing analysis. (F) Bar plot depicting the distribution of cells from different species within a specific subcluster identified in single-cell RNA sequencing analysis. (G) Inferred developmental trajectory of recipient-derived T cells. Arrows in the main plot revealed a dichotomic cell fate division. (H) Trajectory plots displaying levels of cytotoxic and exhaustion scores in recipient-derived T cells, with the expression level of signature genes (KLRD1 for cytotoxicity, PDCD1 for exhaustion). (I) Immunofluorescence staining of CD8+ T cell with signature proteins. Scale bars, 20 µm. (J) Dot plots showing ligand-receptor interactions between recipient-derived cytotoxic T cells and indicated kidney cell subsets. P values were calculated using an empirical shuffling test.

We next interrogated the differentiation state of graft-infiltrating CD8^+^ T cells. Pseudotemporal trajectory analysis inferred a bifurcated path originating from an activated precursor state (**Fig. 8F, Extended Data Fig. 9C-9E, Supplementary Table 12**). One branch (Fate 1) progressed through states co-expressing cytotoxic effectors (GZMA/B, KLRF1) and inhibitory receptors (LAG3, TIGIT), suggesting a cytotoxic state acquiring features of dysfunction. The alternative branch (Fate 2) culminated in a state co-expressing exhaustion markers (PDCD1, CTLA4), the immunomodulatory cytokine IL10, and pro-apoptotic genes, indicating a divergent program blending exhaustion with regulatory potential **(Fig. 8G, 8H, Extended Data Fig. 9F, 9G, Supplementary Table 13)**. This suggests that under rejection pressure, CD8^+^ T cells may adopt heterogeneous functional states beyond a simple effector-exhaustion dichotomy ^35^.

Beyond myeloid interactions, we identified substantial T cell communication with TECs **(Fig. 8I).** The dominant CTL population engaged TECs via both cytotoxic (FASLG-FAS, TNF-TNFRSF1A/B) and inhibitory (LGALS3-MERTK, CD274-CD80) ligand-receptor pairs, pointing to a concurrent balance of attack and suppression. Proliferating T cells signaled to TECs primarily through WNT pathways (WNT8B-FZD1–9), which were co-regulated by secreted inhibitors, implicating this axis in modulating epithelial repair or fibrosis **(Extended Data Fig. 9I)**. Notably, Th2 cells specifically expressed CD40LG, which interacted not only with CD40 on TECs but also with integrin receptors (integrin α5β1 and α5β2). This dual interaction reveals a potential signaling mechanism linking CD40 co-stimulation with integrin-mediated adhesion and matrix remodeling within the rejecting graft **(Extended Data Fig. 9J)**.

## Discussion

Recent studies of pig-to-human xenotransplantation in decedent models have elucidated early immune dynamics and graft adaptation, highlighting human myeloid cell infiltration and activation of tissue-intrinsic repair programs as key features of the xenogeneic response^13,36^. Complementing these clinical observations, we present here a high-resolution single-cell atlas of pig-to-monkey kidney xenotransplantation. Our analysis reveals a dominant innate immune landscape characterized by extensive infiltration of macrophage populations, including recipient-specific subsets, inflammatory priming subpopulations, and functional polarization beyond the classical M1/M2 dichotomy. While consistent with human decedent studies in underscoring the primacy of innate immunity, our primate model uniquely captures the dynamic reciprocity between broad and persistent IFNE expression across multiple tubular cell subsets, which drives recruitment and enrichment of recipient-specific macrophages, and macrophage-mediated intercellular signaling networks within the xenograft kidney; these processes are likely obscured in brain-dead recipients by systemic inflammatory confounders. These findings, enabled by the physiological and immunological integrity of the NHP model, not only provide a basis for deciphering marcrophage-centric pathways that may mediate progressive graft injury and identify several potential interventional targets, but also constitute a high-resolution single-cell dataset that serves as a valuable resource for further understanding xenotransplant immunology and supporting subsequent translational research.

The predominance of recipient-derived macrophages in our atlas highlights a critical, yet unresolved, question in xenotransplantation: which infiltrating subsets are model-specific, and which are fundamental to the xenogeneic response? Our identification of distinct subsets, including FN1^+^, MARCO^+^, IDO1^+^, and ACKR1^+^ macrophages, as exclusively of recipient origin suggests a profound reshaping of the myeloid landscape. Intriguingly, while pig-to-human decedent models also report robust infiltration of human macrophages and inflammatory activation, the resolution of specific transcriptomic subsets defined by markers such as FN1 or ACKR1 has not been equivalently detailed. This may reflect differences in species biology, analytical depth, or the confounding systemic inflammation inherent to the decedent setting.

Among these, MARCO^+^ macrophages engaged xenograft cells mainly via Prostaglandin E2, HBEGF, and TGFB1 to coordinate anti-inflammatory and profibrotic signaling, alongside with significant enrichment of Cholesterol-LIPA-RORA/C interactions. This finding aligns with documented characteristics of cholesterol metabolic dysregulation^37,38^, thereby further confirming the metabolic reprogramming properties of this cellular subset. Therapeutically, this subset’s MARCO receptor offers promise, as nanoparticle targeting redirects such macrophages to spleen for apoptosis, suppressing rejection^39^. ACKR1, as a transmembrane receptor, scavenges chemokines across CXC and CC subfamilies without initiating downstream signaling, thereby effectively clearing these chemo-attractants to mitigate inflammatory responses^40,41^. Notwithstanding the lack of established spatiotemporal correlations between ACKR1⁺ macrophage infiltration and chemokine levels in our study, these cells similarly functioned as anti-inflammatory hubs via PGE2, TGFB1, HBEGF crosstalk with xenografts while suppressing T cells via CD274/PDCD1LG2-PDCD1, CD80/86-CTLA4 and NECTIN2-TIGIT interactions. Collectively, these subsets point to a sophisticated division of labor within the recipient’s myeloid response. They appear not merely as passive infiltrates but as active architects of the graft microenvironment, integrating metabolic reprogramming, matrix remodeling, and immune modulation.

Building upon this cellular atlas, we further identified three key pro-inflammatory macrophage subsets (APOE^+^, CCL2^+^, and IL-1A^+^) that mediate inflammatory injury in xenograft kidneys by evaluating the expression patterns of pro-inflammatory damage mediators, and proposed potential therapeutic agents based on their transcriptional signatures. Computational drug prediction^42,43^ converged not on a single target but on two complementary classes of agents: T cell co-stimulation blockers (belatacept/abatacept) and a myeloid-directed CSF1R inhibitor (pexidartinib). This dual targeting implies that curbing xenograft inflammation may require coordinated intervention across both the adaptive immune axis, centered on antigen presentation and T cell priming, and the innate axis driven by macrophage recruitment and activation. The inclusion of CTLA4-Ig fusion proteins is particularly salient. While their established mechanism involves blocking CD28-mediated T cell co-stimulation via CD80/CD86 on antigen-presenting cells^44,45^, our finding that infiltrating macrophages abundantly express these ligands suggests a broader potential immunomodulatory role. In the xenograft microenvironment, where macrophages may contribute substantially to early inflammation and antigen presentation^46,47^, belatacept may simultaneously dampen myeloid-dependent T cell activation and mitigate downstream donor-specific antibody responses^44^, a dual effect that could be especially advantageous in the highly immunogenic setting of xenotransplantation. Complementing this approach, pexidartinib offers a direct means to blunt the innate inflammatory wave by depleting CSF1R-dependent macrophages^48^. CSF1/CSF1R pathway is essential for monocyte differentiation, survival, proliferation, and functional maintenance, particularly for tissue-resident macrophages; by blocking this signaling, pexidartinib can effectively deplete macrophages in tissues^49^. Its prediction here reinforces the notion that macrophage-driven injury represents a tractable checkpoint in xenograft rejection. Although these agents mechanistically target adaptive and innate immunity respectively and theoretically possess the potential to modulate macrophage-mediated inflammatory injury in xenotransplantation, their efficacy, safety, and optimal timing in this setting remain to be experimentally validated.

Beyond macrophage-centric mechanisms, our study reveals a profound rewiring of the interferon signaling network, characterized by a relative attenuation of the canonical pro-inflammatory axis driven by type II IFN-γ and a concomitant, unexpected prominence of epithelial-immune communication mediated by type I IFN-ε. In transplant immunology, IFN-γ is a central effector cytokine produced by infiltrating effector T cells and natural killer cells^50,51^. It activates graft endothelial cells, parenchymal cells, and resident myeloid populations, inducing the expression of effector molecules such as CXCL9 and CXCL10, thereby fueling inflammation and amplifying immune cell recruitment. This pathway constitutes a molecular cornerstone of both T cell-mediated and antibody-mediated rejection^50,52^. Notably, in our xenotransplant model, despite significant immune infiltration, the response of renal tubular epithelial cells to this canonical IFN-γ-inflammatory signal exhibited attenuation and heterogeneity. This observation suggests that under the extreme immunologic challenge of cross-species transplantation, graft parenchyma may engage specific adaptive mechanisms to partially buffer or reshape this potent pro-inflammatory axis, potentially representing an intrinsic tissue-level adaptation to limit collateral damage.

In stark contrast, the pattern of engagement of IFN-ε, an epithelium-derived immunoregulatory factor, reveals a potential counter-balancing mechanism. Our data demonstrate widespread yet heterogeneous expression of IFN-ε across multiple tubular epithelial subsets in the xenograft kidney, coupled with robust inferred ligand-receptor interactions with clusters of IDO1^+^ macrophages, a population endowed with potent immunosuppressive potential^53,54^. This delineates a local immunoregulatory circuit wherein stress-induced IFN-ε upregulation in tubular epithelium recruits and instructs host-derived IDO1^+^ macrophages to exhaust T-cell function via tryptophan metabolism, thereby fostering an immune-tolerant niche that operates in parallel with, and may antagonize, the systemic inflammation driven by IFN-γ.This mechanism aligns with recent findings that IFN-ε can reprogram the tumor immune microenvironment in ovarian cancer, suppressing myeloid-derived suppressor cells and regulatory T cells^55,56^, collectively underscoring IFN-ε’s conserved role as an immunomodulatory node across disparate pathological states. Building on this, we propose an integrative framework that transcends the traditional binary paradigm of “rejection versus tolerance”. The early fate of a xenograft may be determined by the dynamic confliction and spatial competition between two principal signaling axes: the host-systemic immune attack mediated by IFN-γ and the graft-local immunoregulation orchestrated by IFN-ε.

Further reflecting the complexity of the myeloid compartment, we observed that the hybrid M1/M2 macrophage phenotype transcends the classical polarization dichotomy, representing a functionally integrated state that concurrently activates M1 and M2 transcriptional signatures rather than an intermediate continuum. This plasticity is evolutionarily conserved across physiological and pathological contexts: alveolar macrophages in healthy lung co-express CD80/CD86 (M1 markers) and CD206/CD163 (M2 makers) markers^57^, while endometrial-resident macrophages simultaneously exhibit pro-inflammatory and tolerogenic functions^58^. In chronic inflammatory bowel diseases, adenoviral induction of the pro-lymphangiogenic factor VEGF-C reprograms macrophages to upregulate M2 markers (Ym1, Fizz1, Arg1) while retaining high expression of canonical M1 markers (Cox2, iNOS, CD80), forming a hybrid phenotype critical for bacterial clearance and inflammatory cell mobilization^59^. Crucially, pathological microenvironments drive distinct mechanistic trajectories for hybrid states. In our system, recipient-derived macrophages on Fate 2 originate from metabolically reprogrammed precursors (State 3) that acquire pro-inflammatory traits after adopting oxidative phosphorylation-dependent M2-like activity. Conversely, tissue-resident macrophages on Fate 1 undergo unidirectional transition from an M1-dominant to an M2-dominant. This divergence in pseudotemporal dynamics, metabolic priming versus transcriptional shifting, reveals that hybrid phenotypes emerge through specific regulatory logic, not a universal continuum. Our findings thus redefine macrophage plasticity as a “context-coupled activation” paradigm, where tissue niche dictates the mechanistic basis of co-activation, surpassing both binary and linear polarization models.

We acknowledge several critical limitations of this study. First, the study relies solely on GGTA1 knockout pigs without incorporating clinically relevant multi-gene edits (e.g., human CD46, CD47, and THBD genes), which obscures genotype-specific immune regulatory mechanisms (such as the regulation of macrophage-mediated phagocytosis by CD47). This limits its translational value, as the observed mechanisms may not recapitulate those in advanced xenograft models critical for mitigating hyperacute and acute rejection. Second, the small sample size (n=2 per group) and lack of validation studies (e.g., marker-based cluster confirmation and functional assays) undermine the rigor, reproducibility, and generalizability of the results, precluding definitive differentiation between circulating and infiltrative immune signatures as well as verification of rejection-specific responses. Third, using non-transplanted porcine kidneys as controls introduces significant confounding factors, as these controls have not undergone ischemia-reperfusion injury, transplantation procedures, or immunosuppressive treatment, hindering the precise distinction between rejection-driven changes and baseline xenogeneic variability. Finally, the short graft survival times (7 and 11 days) prevent the analysis of long-term immune adaptation or chronic rejection mechanisms; meanwhile, the absence of key data related to primary graft outcomes, such as renal function evaluation and urine output from the transplanted kidneys, hinders the rational interpretation of histological findings and single-cell RNA sequencing data within a clinical context. These limitations constrain the translational application of our findings; future studies should employ multi-gene-edited models, expand sample sizes, establish validated analytical pipelines, set up matched controls, and extend follow-up durations to thoroughly explore mechanisms underlying long-term graft survival and dynamic immune responses.

In summary, this study establishes the first high-resolution single-cell atlas of pig-to-NHP kidney xenotransplantation. Our analysis reveals a dominant innate immune landscape characterized by extensive macrophage infiltration and activation, and identifies multiple macrophage subsets with distinct functions and origins. Furthermore, we uncover an immunoregulatory axis driven by graft-derived IFN-ε, a finding that provides a novel perspective on intrinsic immunomodulatory mechanisms within the xenograft. Although this study has limitations regarding the genetic engineering model, sample size, and observational duration, our findings nevertheless establish a molecular framework for understanding xenotransplant immunodynamics and highlight potential therapeutic strategies. Thus, this work not only provides a key resource for deciphering macrophage-mediated injury pathways but also lays an important foundation for refining immunosuppressive regimens and engineering donor organs to advance the clinical translation of xenotransplantation.

## Methods

### Ethics acquisition

All animal experiments were conducted in accordance with protocols jointly reviewed and approved by the Institutional Animal Ethics Committees of Chinese PLA General Hospital and Fuwai Hospital (Approval No. FWAEC-JL-010-1/0-2020). Following institutional approval, the final application was submitted for regulatory compliance verification to relevant government authorities including the Beijing Municipal Science and Technology Commission and Beijing Municipal Health Commission, as well as academic oversight bodies comprising the Chinese Academy of Medical Sciences and Medical School of Chinese PLA.

### Bio-safety standards and management

The entire xenotransplantation protocol, encompassing animal husbandry, porcine kidney procurement, transplantation surgery, postoperative care, and scientific observation, was conducted within the biosafety-controlled environment of Beijing key laboratory of preclinical research and evaluation for cardiovascular implant materials. This facility maintains research infrastructure compliant with Category B infectious disease containment standards as stipulated by China’s Law on Prevention and Treatment of Infectious Diseases. All biological materials including non-human primate and porcine remains, along with associated medical waste, underwent high-temperature incineration through a closed-loop disposal system to ensure complete biological decontamination.

### Selection and characterization of gene-edited porcine donors and recipients

Two GGTA1-knockout Bama miniature pigs were generated by Clonorgan Biotechnology Co., Ltd (Chengdu, China)^60^ and housed under designated pathogen-free (DPF) conditions. Two rhesus macaques (Macaca mulatta) were provided by Beijing Prima Biotech Inc (Beijing, China). The donor pigs and recipient monkeys underwent climate-controlled ground transportation with biosecurity protocols ensuring complete isolation during transfer from Chengdu to the specific pathogen-free (SPF) facility at Beijing key laboratory of preclinical research and evaluation for cardiovascular implant materials.

Donor 1: Male (68 days old, 6.5 kg, blood type O) with monoallelic GGTA1 knockout (GTKO) confirmed by genomic sequencing.

Donor 2: Male (67 days old, 6.5 kg, blood type O) carrying identical monoallelic GGTA1 knockout (GTKO) configuration.

Recipient 1: Male, 10 years old, 9 kg, blood type O.

Recipient 2: Male, 11 years old, 9.5 kg, blood type O.

### Surgery

Donor porcine kidneys were procured under standardized general anesthesia using open surgical techniques mirroring clinical living donor nephrectomy protocols. Following in situ perfusion with 4°C histidine-tryptophan-ketoglutarate (HTK) solution, the organs were preserved in cold HTK and transported to the recipient operating suite. In parallel, recipient monkeys underwent general anesthesia and midline laparotomy. Through a right retroperitoneal approach, the iliac fossa was exposed by peritoneal reflection. Vascular anastomoses were performed using end-to-side techniques: (1) Renal artery to external iliac artery; (2) Renal vein to external iliac vein; (3) Ureteral continuity was established via direct implantation into the bladder dome, supplemented by double-J stent placement within the renal pelvis. Immediate graft reperfusion was confirmed following vascular anastomosis completion.

### Immunosuppression regimens

Both xenograft recipients received a dual-phase immunosuppressive regimen mirroring clinical allotransplantation protocols, comprising induction and maintenance phases.

Phase 1: induction Therapy

Initiated preoperatively and administered as: IL-2 receptor antagonist (Basiliximab): 20 mg/dose IV preoperatively and on postoperative day 1-4; Methylprednisolone: 500 mg IV bolus preoperatively and post-reperfusion; Anti-CD20 therapy (Rituximab): Single intraoperative dose (20 mg/kg).

Phase 2: Maintenance Therapy

Postoperative management included: Mycophenolate mofetil: 1.5 mg/12h PO; Corticosteroid taper: Initiated at 1 mg/kg/day IV, reduced by 10 mg every; 72 hr to 10 mg/day maintenance; Cyclosporine A: Titrated to target trough concentration (0.20-0.35 μg/mL) via therapeutic drug monitoring.

### Experimental endpoints

The preset endpoint was irreversible graft dysfunction unless one of the following scenarios occurred: heart-lung dysfunction, uncontrollable infections, or termination requested by the ethics committee.

### Histological evaluation and Multiplexed Immunofluorescence

Xenotransplanted kidney specimens harvested at experimental endpoints were fixed in 10% neutral-buffered formalin (Fisher Scientific) for 48 hours, followed by standardized tissue dehydration and paraffin embedding using automated processors. Serial 5-μm sections were systematically prepared for Hematoxylin & eosin, immunohistochemistry (IHC) and multiplexed immunofluorescence staining (IF). The primary antibody panel including: anti-IgM (11016-1-AP, 1:100, Proteintech), anti-IgG (ab109489, 1:500, Abcam), C4d (22233-1-AP, 1:200, Proteintech), anti-CD31 (ab28364, 1:50, Abcam), anti-vWF (ab6994, 1:200, Abcam), anti-CD3 (GB12014, 1:1000, Servicebio) and anti-CD15 (YT0726, 1:200, Immunoway) for IHC; anti-CD68 (bs-0649R, 1:200, Bioss), anti-SPP1 (83341-1-RR, 1:200, Proteintech), anti-VEGFA (66828-1-Ig, 1:200, Proteintech), anti-PD-L1 (ab205921, 1:200, Abcam), anti-IDO1 (ab211017, 1:1000, Abcam), anti-ARG1 (66129-1-Ig, 1:200, Proteintech), anti-FCGR2B (NB100-65338, 1:100, Novus Biologicals), anti-CD8a (66868-1-Ig, 1:400, Proteintech), and anti-KLRD1 (84466-5-RR, 1:200, Proteintech) for multiplexed IF.

### Tissue processing and single cell processing

Fresh xenograft and control kidney samples were washed three times with phosphate-buffered saline (PBS) and minced into small pieces (<1 mm^3^) on ice. Single cells from the kidney were isolated using the Multi Tissue Dissociation Kit as protocol (130-110-201, Miltenyi Biotec, Germany). After digestion, samples were filtered using a 70 mm nylon strainer (Thermo Fisher Scientific), washed with 1% bovine serum albumin (BSA, Sigma-Aldrich) and 2 mM EDTA in PBS. Optionally, red blood cells in single-cell suspensions were removed using the Red Blood Cell Lysis Solution (10×) Kit (Miltenyi Biotec), according to the manufacturer’s specifications. The cell pellet was resuspended in PBS containing 1% BSA. The cells were stained with live/dead cell dyes (559925, BD Pharmingen™ 7-AAD) and anti-CD45 antibodies (561865, BD Pharmingen™ FITC Mouse Anti-Human CD45). Live kidney single cells and CD45^+^ immune cells were sorted using flow cytometry (BD FACS Aria II) for the construction of the single-cell library. Only suspensions with a viability of ≥85% and cell agglomeration rates of ≤10% were regarded as qualified samples for library construction. Cells of qualified samples were then diluted with PBS containing 1% BSA to about 700–1,200 cells per µl for scRNA-seq. Single-cell suspensions were processed using the Chromium Next GEM Single Cell 3′ GEM, Library & Gel Bead Kit v.3.1 (10× Genomics) and run in the Chromium Controller following the manufacturer’s protocol, with a capture target of 16, 500 cells for each sample. A chromium single-cell 5′ library was then constructed with Chromium Next GEM Single Cell 3′ Library Construction Kit v.3.1 (10× Genomics) with quality-qualified cDNA. After fragmentation, adaptor ligation and sample index PCR, the library was finally quantitatively examined. The final library pool was sequenced on an Illumina NovaSeq 6000 instrument using 150 base pair paired-end reads.

### Data pre-processing and quality control of scRNA-seq

For xenotransplantation samples, we downloaded reference genomes and GTF annotation files from NCBI, specifically GCF-003339765.1_Mmul10 and GCF_000003025.6_Sscrofa11.1. A combined reference genome was constructed following the multi-species reference genome guidelines provided by 10x Genomics (https://support.10xgenomics.com/single-cell-gene-expression/software/pipelines/latest/advanced/references#multi). Non-xenotransplantation samples were analyzed using the Sscrofa11.1 reference genome. Raw single-cell sequencing data were processed with the CellRanger (v8.0.1) pipeline with default parameters to generate single-cell count matrix and fed to Seurat (v4.3.0) for downstream analysis. DecontX (v1.6.0) was used to estimate and remove ambient RNA contamination, we employed the Scrublet (v0.2.3) algorithm to identify and exclude potential doublets from our single cell data. Cells were retained based on these thresholds: total counts below 15,000, more than 200 detected genes, mitochondrial gene content below 5%, ribosomal gene content below 50%, and Scrublet doublet scores less than 0.3. applying above threashold, a total of 75,324 high quality cells were kept for further analysis. Expression values were then scaled to 10,000 transcripts per cell and Log-transformed. Effects of variable (percent. mito) were estimated and regressed out using a GLM (ScaleData function, model.use = ‘linear’), and the scaled and centered residuals were used for dimensionality reduction and clustering.

### Dimension reduction and unsupervised clustering

Top 2,000 Highly variable genes were generated using Seurat ‘vst’ option and used for subsequent principal component analysis and dimensional reduction. To reduce dimensionality of the datasets, the RunPCA function was conducted with default parameters on linear-transformation scaled data generated by the ScaleData function.. Batch correction between different samples was achieved using Harmony (v1.2.1). The first 30 principal components (PCs) were utilized to construct a shared nearest neighbor (SNN) graph via the FindNeighbors function. Subsequently, cell clustering was performed using the FindClusters function, implementing the Louvain algorithm. Finally, we performed non-linear dimensional reduction with the RunUMAP function with default settings.

### Cell type Annotation

DEGs of a certain cell cluster were identified compared to all other cell clusters using Wilcox rank-sum test followed by Bonferroin correction by Seurat option ‘FindAllMarkers’. Only upregulated DEGs within each cluster were considered. Major cell types were assigned to Seurat clusters according to the normalized expression of canonical markers. The kidney cells and immune cell groups, excluding LRRC43^+^ and RHOH^+^ unknown cell groups, were further re-clustered to reveal subtypes within major cell clusters. By repeating the above steps mentioned (normalization, dimensional reduction, batch effect correction and clustering), different specific cell subtypes were identified and annotated according to the average expression of canonical marker genes. For each subcluster of a major cell type. The selection criteria of the marker gene for annotation included (1) top-ranking DEGs for the corresponding cell cluster; (2) strong specificity of gene expression, meaning a high expression ratio within the corresponding cell cluster but low in other clusters; and (3) literature supports that it is either a marker gene or functionally relevant to the type of cell. Markers used in this pipeline are addressed in Supplementary Table 2.

### Functional enrichment analysis

To characterize the biological implications of the DEGs, we performed Gene Ontology (GO) functional annotation and Kyoto Encyclopedia of Genes and Genomes (KEGG) pathway enrichment analysis using the R package clusterProfiler (v4.10.1). GO analysis encompassed three categories: Biological Process (BP), Molecular Function (MF), and Cellular Component (CC). The statistical significance of enriched terms was evaluated using a hypergeometric test, the BenjaminiHochberg method was used to estimate the false discovery rate (FDR). Terms with an adjusted P-value < 0.05 and a raw P-value < 0.01 were considered significantly enriched. The results were visualized as bubble plots or bar charts to highlight the most prominent biological signatures.

### Trajectory analysis

To delineate the dynamic transition states of Macrophages and T cells, we performed pseudotime trajectory inference using the Monocle2 R package (v2.30.1). The raw count matrices and associated metadata were exported from the Seurat object and initialized as a CellDataSet object via the import CDS function. To identify the developmental progression, we employed a data-driven approach to select highly variable genes (HVGs) for cell ordering, ensuring that the trajectory captured the most biologically relevant transcriptional changes. The high-dimensional gene expression data were then projected into a lower-dimensional space using the Discriminative Dimensional Reduction Tree (DDRTree) algorithm. The minimum spanning tree was constructed with the reduceDimension function (parameters: max_components = 2, num_dim = 10, norm_method = “log”, and scaling = TRUE). Finally, cells were ordered along the trajectory branches, and pseudotime was calculated to represent the relative progression of Macrophage and T cell differentiation.

### Cellular interaction analysis

We applied the CellphoneDB (v5.0.0) of known receptor-ligand pairs to assess cell–cell communication in our dataset. The normalized gene expression matrix was converted into the required format: a gene expression table (genes × cells) and a corresponding metadata table containing cell barcode and their assigned labels. Parameters included 1,000 permutations for robust P-value estimation and a minimum expression threshold of 0.1. Interactions were trimmed based on significant sites with P-value < 0.01. The relative expression levels (Z-scores) and P-value of ligand–receptor molecule pairs are displayed in the form of dot plots or interaction heatmaps.

### Transcription factor activity analysis

We used pySCENIC (v0.12.1) to infer transcription factors. The analysis followed a three-step workflow: First, potential TF-target co-expression modules were identified using GENIE3, based on the regression of TF expression levels against potential targets. Second, these modules were refined using RcisTarget to retain only those with significant TF-binding motif enrichment, thereby pruning indirect targets and forming high-confidence regulons. Finally, the activity of each regulon within individual cells was quantified using the AUCell algorithm. Regulons with highest activity in each group were identified by Wilcox rank-sum test followed by Bonferroni correction with adjusted P-value less than 0.01.

### Scoring gene sets in scRNA-seq data

Selected signatures were scored by the AddModuleScore algorithm in Seurat using the gene sets listed in Supplementary Table 3. All parameters were set to the default.

### Drug prediction

Candidate drugs were extracted from ChEMBL ^42^ database under ATC/DDD classification L, excluding L03 which is immunostimulants. Resulting of 257 candidate drugs. drug2cell (v0.1.2) ^43^ function ‘score’ was utilized to score each drug. Using drug score as input, we performed Wilcox rank-sum test followed by Bonferroni correction between our target cell group and the remaining cell group, a cutoff of avg_log2FC > 0.25 and adjust P-value < 0.01 was set to look for potential drugs.

### Statistics and reproducibility

Wilcox rank-sum test was used for comparison between two groups. Multiple comparison corrections were applied using Bonferroni method, ensuring robustness of the findings. Statistical significance was defined as P-value < 0.01 unless otherwise indicated.

## Data availability

Processed single-cell RNA-seq data (cellranger) of this study have been deposited in the OMIX, China National Center for Bioinformation/Beijing Institute of Genomics, Chinese Academy of Sciences (https://ngdc.cncb.ac.cn/omix: accession no.OMIX011460). The data that support the findings of this study are also available from the corresponding author upon request.

## Code availability

Codes used in this study are available at BioCode (https://ngdc.cncb.ac.cn/biocode/tool/7990). No custom code was developed for this study.

## Acknowledgements

The authors thank prof. Jiangping Song, Xiumeng Hua, Yuan Chang and the staff members at the Beijing key laboratory of preclinical research and evaluation for cardiovascular implant materials for their irreplaceable contributions to this animal experiment. We also appreciate Shengyang Wu, Kai Feng and Mingyan Yang for their technical assistance and expertise in transcriptomic data analyses. We also acknowledge the research teams who made their datasets publicly available and thereby enabled cross-cohort validation in this study. This work was sponsored by Frontier Biotechnology Key Project of National Key R & D Program of the Ministry of Science and Technology of China (2023YFC3404300; to Jiangping Song).

## Author Contributions Statement

H.W., J.C., Y.C., X.H., J.P., X.Z., and Y.F. conceived and designed the study. W.C., F.Y. and S.Y. developed the methodology. H.W. J.C., Y.C., X.H., J.P., X.Z., and Y.F. performed the surgery of transplantation. F.Y. and S.Y. carried out investigation and data collection. H.W., J.C., Y.C., C.W., and Y.F. curated and organized the data. H.W. and C.W performed bioinformatics analysis and data visualization. H.W., J.C., and Y.C., drafted the manuscript. W.C., J.S., X.Z., and Y.F. reviewed and edited the manuscript. J.S. acquired funding and provided study resources. W.C., J.S., X.Z., and Y.F. supervised the study. All authors reviewed and approved the final manuscript.

## Reference

1. Matas, A.J., Montgomery, R.A., and Schold, J.D. (2023). The Organ Shortage Continues to Be a Crisis for Patients With End-stage Kidney Disease. JAMA Surg 158, 787–788. 10.1001/jamasurg.2023.0526.

2. Sykes, M., and Sachs, D.H. (2022). Progress in xenotransplantation: overcoming immune barriers. Nat Rev Nephrol 18, 745–761. 10.1038/s41581-022-00624-6.

3. Montgomery, R.A., Mehta, S.A., Parent, B., and Griesemer, A. (2022). Next steps for the xenotransplantation of pig organs into humans. Nat Med 28, 1533–1536. 10.1038/s41591-022-01896-y.

4. Xing, K., Chang, Y., Jia, H., and Song, J. (2025). Advances in Subclinical and Clinical Trials and Immunosuppressive Therapies in Xenotransplantation. Xenotransplantation 32, e70053. 10.1111/xen.70053.

5. Phelps, C.J., Koike, C., Vaught, T.D., Boone, J., Wells, K.D., Chen, S.H., Ball, S., Specht, S.M., Polejaeva, I.A., Monahan, J.A., et al. (2003). Production of alpha 1,3-galactosyltransferase-deficient pigs. Science 299, 411–414. 10.1126/science.1078942.

6. Wijkstrom, M., Iwase, H., Paris, W., Hara, H., Ezzelarab, M., and Cooper, D.K. (2017). Renal xenotransplantation: experimental progress and clinical prospects. Kidney Int 91, 790–796. 10.1016/j.kint.2016.08.035.

7. Cooper, D.K.C., and Pierson, R.N., 3rd (2023). Milestones on the path to clinical pig organ xenotransplantation. Am J Transplant 23, 326–335. 10.1016/j.ajt.2022.12.023.

8. Schmauch, E., Piening, B.D., Dowdell, A.K., Mohebnasab, M., Williams, S.H., Stukalov, A., Robinson, F.L., Bombardi, R., Jaffe, I., Khalil, K., et al. (2025). Multi-omics analysis of a pig-to-human decedent kidney xenotransplant. Nature. 10.1038/s41586-025-09846-7.

9. Wang, Y., Chen, G., Pan, D., Guo, H., Jiang, H., Wang, J., Feng, H., He, S., Du, J., Zhang, M., et al. (2024). Pig-to-human kidney xenotransplants using genetically modified minipigs. Cell Rep Med 5, 101744. 10.1016/j.xcrm.2024.101744.

10. Montgomery, R.A., Stern, J.M., Lonze, B.E., Tatapudi, V.S., Mangiola, M., Wu, M., Weldon, E., Lawson, N., Deterville, C., Dieter, R.A., et al. (2022). Results of Two Cases of Pig-to-Human Kidney Xenotransplantation. N Engl J Med 386, 1889–1898. 10.1056/NEJMoa2120238.

11. Judd, E., Kumar, V., Porrett, P.M., Hyndman, K.A., Anderson, D.J., Jones-Carr, M.E., Shunk, A., Epstein, D.R., Fatima, H., Katsurada, A., et al. (2024). Physiologic homeostasis after pig-to-human kidney xenotransplantation. Kidney Int 105, 971–979. 10.1016/j.kint.2024.01.016.

12. Loupy, A., Goutaudier, V., Giarraputo, A., Mezine, F., Morgand, E., Robin, B., Khalil, K., Mehta, S., Keating, B., Dandro, A., et al. (2023). Immune response after pig-to-human kidney xenotransplantation: a multimodal phenotyping study. Lancet 402, 1158–1169. 10.1016/S0140-6736(23)01349-1.

13. Pan, W., Zhang, W., Zheng, B., Camellato, B.R., Stern, J., Lin, Z., Khodadadi-Jamayran, A., Kim, J., Sommer, P., Khalil, K., et al. (2024). Cellular dynamics in pig-to-human kidney xenotransplantation. Med 5, 1016–1029 e1014. 10.1016/j.medj.2024.05.003.

14. Montgomery, R.A., Stern, J.M., Fathi, F., Suek, N., Kim, J.I., Khalil, K., Vermette, B., Tatapudi, V.S., Mattoo, A., Skolnik, E.Y., et al. (2025). Physiology and immunology of a pig-to-human decedent kidney xenotransplant. Nature 10.1038/s41586-025-09847-6.

15. Du, X., Chang, Y., and Song, J. (2025). Use of Brain Death Recipients in Xenotransplantation: A Double-Edged Sword. Xenotransplantation 32, e70010. 10.1111/xen.70010.

16. Meyfroidt, G., Gunst, J., Martin-Loeches, I., Smith, M., Robba, C., Taccone, F.S., and Citerio, G. (2019). Management of the brain-dead donor in the ICU: general and specific therapy to improve transplantable organ quality. Intensive Care Med 45, 343–353. 10.1007/s00134-019-05551-y.

17. Ganchiku, Y., and Riella, L.V. (2022). Pig-to-human kidney transplantation using brain-dead donors as recipients: One giant leap, or only one small step for transplantkind? Xenotransplantation 29, e12748. 10.1111/xen.12748.

18. Adams, A.B., Kim, S.C., Martens, G.R., Ladowski, J.M., Estrada, J.L., Reyes, L.M., Breeden, C., Stephenson, A., Eckhoff, D.E., Tector, M., and Tector, A.J. (2018). Xenoantigen Deletion and Chemical Immunosuppression Can Prolong Renal Xenograft Survival. Ann Surg 268, 564–573. 10.1097/SLA.0000000000002977.

19. Yamada, K., Yazawa, K., Shimizu, A., Iwanaga, T., Hisashi, Y., Nuhn, M., O’Malley, P., Nobori, S., Vagefi, P.A., Patience, C., et al. (2005). Marked prolongation of porcine renal xenograft survival in baboons through the use of alpha1,3-galactosyltransferase gene-knockout donors and the cotransplantation of vascularized thymic tissue. Nat Med 11, 32–34. 10.1038/nm1172.

20. Kim, S.C., Mathews, D.V., Breeden, C.P., Higginbotham, L.B., Ladowski, J., Martens, G., Stephenson, A., Farris, A.B., Strobert, E.A., Jenkins, J., et al. (2019). Long-term survival of pig-to-rhesus macaque renal xenografts is dependent on CD4 T cell depletion. Am J Transplant 19, 2174–2185. 10.1111/ajt.15329.

21. Hu, X., Tediashvili, G., Gravina, A., Stoddard, J., McGill, T.J., Connolly, A.J., Deuse, T., and Schrepfer, S. (2025). Inhibition of polymorphonuclear cells averts cytotoxicity against hypoimmune cells in xenotransplantation. Nat Commun 16, 3706. 10.1038/s41467-025-58774-7.

22. Cooper, D.K., Ekser, B., Ramsoondar, J., Phelps, C., and Ayares, D. (2016). The role of genetically engineered pigs in xenotransplantation research. J Pathol 238, 288–299. 10.1002/path.4635.

23. Bourke, N.M., Achilles, S.L., Huang, S.U., Cumming, H.E., Lim, S.S., Papageorgiou, I., Gearing, L.J., Chapman, R., Thakore, S., Mangan, N.E., et al. (2022). Spatiotemporal regulation of human IFN-epsilon and innate immunity in the female reproductive tract. JCI Insight 7. 10.1172/jci.insight.135407.

24. Yang, J., Song, X., Zhang, H., Liu, Q., Wei, R., Guo, L., Yuan, C., Chen, F., Xue, K., Lai, Y., et al. (2024). Single-cell transcriptomic landscape deciphers olfactory neuroblastoma subtypes and intra-tumoral heterogeneity. Nat Cancer 5, 1919–1939. 10.1038/s43018-024-00855-5.

25. Vergadi, E., Ieronymaki, E., Lyroni, K., Vaporidi, K., and Tsatsanis, C. (2017). Akt Signaling Pathway in Macrophage Activation and M1/M2 Polarization. J Immunol 198, 1006–1014. 10.4049/jimmunol.1601515.

26. Griffiths, H.R., Gao, D., and Pararasa, C. (2017). Redox regulation in metabolic programming and inflammation. Redox Biol 12, 50–57. 10.1016/j.redox.2017.01.023.

27. Sun, H.J., Zhou, T., Wang, Y., Fu, Y.W., Jiang, Y.P., Zhang, L.H., Zhang, C.B., Zhou, H.L., Gao, B.S., Shi, Y.A., and Wu, S. (2011). Macrophages and T lymphocytes are the predominant cells in intimal arteritis of resected renal allografts undergoing acute rejection. Transpl Immunol 25, 42–48. 10.1016/j.trim.2011.04.002.

28. Mannon, R.B. (2012). Macrophages: contributors to allograft dysfunction, repair, or innocent bystanders? Curr Opin Organ Transplant 17, 20–25. 10.1097/MOT.0b013e32834ee5b6.

29. Dangi, A., Natesh, N.R., Husain, I., Ji, Z., Barisoni, L., Kwun, J., Shen, X., Thorp, E.B., and Luo, X. (2020). Single cell transcriptomics of mouse kidney transplants reveals a myeloid cell pathway for transplant rejection. JCI Insight 5. 10.1172/jci.insight.141321.

30. Ross, G.D. (2000). Regulation of the adhesion versus cytotoxic functions of the Mac-1/CR3/alphaMbeta2-integrin glycoprotein. Crit Rev Immunol 20, 197–222.

31. Xiao, L., Zhang, Z., Luo, X., Yang, H., Li, F., and Wang, N. (2016). Retinoid acid receptor-related orphan receptor alpha (RORalpha) regulates macrophage M2 polarization via activation of AMPKalpha. Mol Immunol 80, 17–23. 10.1016/j.molimm.2016.10.006.

32. Webb, L.M., Sengupta, S., Edell, C., Piedra-Quintero, Z.L., Amici, S.A., Miranda, J.N., Bevins, M., Kennemer, A., Laliotis, G., Tsichlis, P.N., and Guerau-de-Arellano, M. (2020). Protein arginine methyltransferase 5 promotes cholesterol biosynthesis-mediated Th17 responses and autoimmunity. J Clin Invest 130, 1683–1698. 10.1172/JCI131254.

33. Huang, D., Chen, X., Zeng, X., Lao, L., Li, J., Xing, Y., Lu, Y., Ouyang, Q., Chen, J., Yang, L., et al. (2021). Targeting regulator of G protein signaling 1 in tumor-specific T cells enhances their trafficking to breast cancer. Nat Immunol 22, 865–879. 10.1038/s41590-021-00939-9.

34. Stark, J.M., Tibbitt, C.A., and Coquet, J.M. (2019). The Metabolic Requirements of Th2 Cell Differentiation. Front Immunol 10, 2318. 10.3389/fimmu.2019.02318.

35. Zheng, C., Zheng, L., Yoo, J.K., Guo, H., Zhang, Y., Guo, X., Kang, B., Hu, R., Huang, J.Y., Zhang, Q., et al. (2017). Landscape of Infiltrating T Cells in Liver Cancer Revealed by Single-Cell Sequencing. Cell 169, 1342–1356 e1316. 10.1016/j.cell.2017.05.035.

36. Ribas, G.T., Cunha, A.F., Avila, J.P., Giarraputo, A., Morena, L., Lima, K., Gassen, R.B., Chen, J.Y., Lin, J.R., Santagata, S., et al. (2026). Immune profiling in a living human recipient of a gene-edited pig kidney. Nat Med 32, 270–280. 10.1038/s41591-025-04053-3.

37. Brunner, J.S., Vogel, A., Lercher, A., Caldera, M., Korosec, A., Puhringer, M., Hofmann, M., Hajto, A., Kieler, M., Garrido, L.Q., et al. (2020). The PI3K pathway preserves metabolic health through MARCO-dependent lipid uptake by adipose tissue macrophages. Nat Metab 2, 1427–1442. 10.1038/s42255-020-00311-5.

38. Zhang, Y., Yang, W., Kumagai, Y., Loza, M., Yang, Y., Park, S.J., and Nakai, K. (2024). In Silico Analysis Revealed Marco (SR-A6) and Abca1/2 as Potential Regulators of Lipid Metabolism in M1 Macrophage Hysteresis. Int J Mol Sci 26. 10.3390/ijms26010111.

39. Zhou, G., Zhang, L., and Shao, S. (2024). The application of MARCO for immune regulation and treatment. Mol Biol Rep 51, 246. 10.1007/s11033-023-09201-x.

40. Mange, K.C., Prak, E.L., Kamoun, M., Du, Y., Goodman, N., Danoff, T., Hoy, T., Newman, M., Joffe, M.M., and Feldman, H.I. (2004). Duffy antigen receptor and genetic susceptibility of African Americans to acute rejection and delayed function. Kidney Int 66, 1187–1192. 10.1111/j.1523-1755.2004.00871.x.

41. Dawson, T.C., Lentsch, A.B., Wang, Z., Cowhig, J.E., Rot, A., Maeda, N., and Peiper, S.C. (2000). Exaggerated response to endotoxin in mice lacking the Duffy antigen/receptor for chemokines (DARC). Blood 96, 1681–1684.

42. Gaulton, A., Bellis, L.J., Bento, A.P., Chambers, J., Davies, M., Hersey, A., Light, Y., McGlinchey, S., Michalovich, D., Al-Lazikani, B., and Overington, J.P. (2012). ChEMBL: a large-scale bioactivity database for drug discovery. Nucleic Acids Res 40, D1100–1107. 10.1093/nar/gkr777.

43. Kanemaru, K., Cranley, J., Muraro, D., Miranda, A.M.A., Ho, S.Y., Wilbrey-Clark, A., Patrick Pett, J., Polanski, K., Richardson, L., Litvinukova, M., et al. (2023). Spatially resolved multiomics of human cardiac niches. Nature 619, 801–810. 10.1038/s41586-023-06311-1.

44. Badell, I.R., Karadkhele, G.M., Vasanth, P., Farris, A.B., 3rd, Robertson, J.M., and Larsen, C.P. (2019). Abatacept as rescue immunosuppression after calcineurin inhibitor treatment failure in renal transplantation. Am J Transplant 19, 2342–2349. 10.1111/ajt.15319.

45. Leibler, C., Matignon, M., Moktefi, A., Samson, C., Zarour, A., Malard, S., Boutin, E., Pilon, C., Salomon, L., Natella, P.A., et al. (2019). Belatacept in renal transplant recipient with mild immunologic risk factor: A pilot prospective study (BELACOR). Am J Transplant 19, 894–906. 10.1111/ajt.15229.

46. Malone, A.F., Wu, H., Fronick, C., Fulton, R., Gaut, J.P., and Humphreys, B.D. (2020). Harnessing Expressed Single Nucleotide Variation and Single Cell RNA Sequencing To Define Immune Cell Chimerism in the Rejecting Kidney Transplant. J Am Soc Nephrol 31, 1977–1986. 10.1681/ASN.2020030326.

47. Shen, Q., Zhu, T., Yan, S., Wang, Y., Wang, C., Teng, L., Feng, S., Huang, X.R., Tang, S.C.W., Huang, H., et al. (2025). Macrophages mediate acute kidney allograft rejection via a toll-like receptor 4-dependent mechanism. Kidney Int 108, 1105–1122. 10.1016/j.kint.2025.09.014.

48. Lamb, Y.N. (2019). Pexidartinib: First Approval. Drugs 79, 1805–1812. 10.1007/s40265-019-01210-0.

49. Wen, J., Wang, S., Guo, R., and Liu, D. (2023). CSF1R inhibitors are emerging immunotherapeutic drugs for cancer treatment. Eur J Med Chem 245, 114884. 10.1016/j.ejmech.2022.114884.

50. Halloran, P.F., Venner, J.M., Madill-Thomsen, K.S., Einecke, G., Parkes, M.D., Hidalgo, L.G., and Famulski, K.S. (2018). Review: The transcripts associated with organ allograft rejection. Am J Transplant 18, 785–795. 10.1111/ajt.14600.

51. Park, E.M., Lee, H., Kang, H.J., Oh, K.B., Kim, J.S., Chee, H.K., Park, J.H., Park, K.S., and Yun, I.J. (2021). Early Interferon-Gamma Response in Nonhuman Primate Recipients of Solid-Organ Xenotransplantation. Transplant Proc 53, 3093–3100. 10.1016/j.transproceed.2021.09.028.

52. Galdina, V., Puga Yung, G.L., and Seebach, J.D. (2025). Cytotoxic Responses Mediated by NK Cells and Cytotoxic T Lymphocytes in Xenotransplantation. Transpl Int 38, 13867. 10.3389/ti.2025.13867.

53. Uyttenhove, C., Pilotte, L., Theate, I., Stroobant, V., Colau, D., Parmentier, N., Boon, T., and Van den Eynde, B.J. (2003). Evidence for a tumoral immune resistance mechanism based on tryptophan degradation by indoleamine 2,3-dioxygenase. Nat Med 9, 1269–1274. 10.1038/nm934.

54. Brandacher, G., Perathoner, A., Ladurner, R., Schneeberger, S., Obrist, P., Winkler, C., Werner, E.R., Werner-Felmayer, G., Weiss, H.G., Gobel, G., et al. (2006). Prognostic value of indoleamine 2,3-dioxygenase expression in colorectal cancer: effect on tumor-infiltrating T cells. Clin Cancer Res 12, 1144–1151. 10.1158/1078-0432.CCR-05-1966.

55. Marks, Z.R.C., Campbell, N.K., Mangan, N.E., Vandenberg, C.J., Gearing, L.J., Matthews, A.Y., Gould, J.A., Tate, M.D., Wray-McCann, G., Ying, L., et al. (2023). Interferon-epsilon is a tumour suppressor and restricts ovarian cancer. Nature 620, 1063–1070. 10.1038/s41586-023-06421-w.

56. Elorbany, S., Malacrida, B., and Balkwill, F. (2023). Interferon epsilon and ovarian cancer. Trends Cancer 9, 985–986. 10.1016/j.trecan.2023.09.008.

57. Mitsi, E., Kamng’ona, R., Rylance, J., Solorzano, C., Jesus Reine, J., Mwandumba, H.C., Ferreira, D.M., and Jambo, K.C. (2018). Human alveolar macrophages predominately express combined classical M1 and M2 surface markers in steady state. Respir Res 19, 66. 10.1186/s12931-018-0777-0.

58. Quillay, H., El Costa, H., Marlin, R., Duriez, M., Cannou, C., Chretien, F., Fernandez, H., Lebreton, A., Ighil, J., Schwartz, O., et al. (2015). Distinct characteristics of endometrial and decidual macrophages and regulation of their permissivity to HIV-1 infection by SAMHD1. J Virol 89, 1329–1339. 10.1128/JVI.01730-14.

59. D’Alessio, S., Correale, C., Tacconi, C., Gandelli, A., Pietrogrande, G., Vetrano, S., Genua, M., Arena, V., Spinelli, A., Peyrin-Biroulet, L., et al. (2014). VEGF-C-dependent stimulation of lymphatic function ameliorates experimental inflammatory bowel disease. J Clin Invest 124, 3863–3878. 10.1172/JCI72189.

60. Xing, K., Chang, Y., Zhang, X., Du, X., and Song, J. (2025). Xenotransplantation in China: Past, Present, and Future. Xenotransplantation 32, e70038. 10.1111/xen.70038.

