## Extended Data Figures 1-9 for "Single-cell atlas of pig-to-monkey kidney xenotransplantation reveals macrophage chimerism and an IFN-ε orchestrated graft protective immune niche"

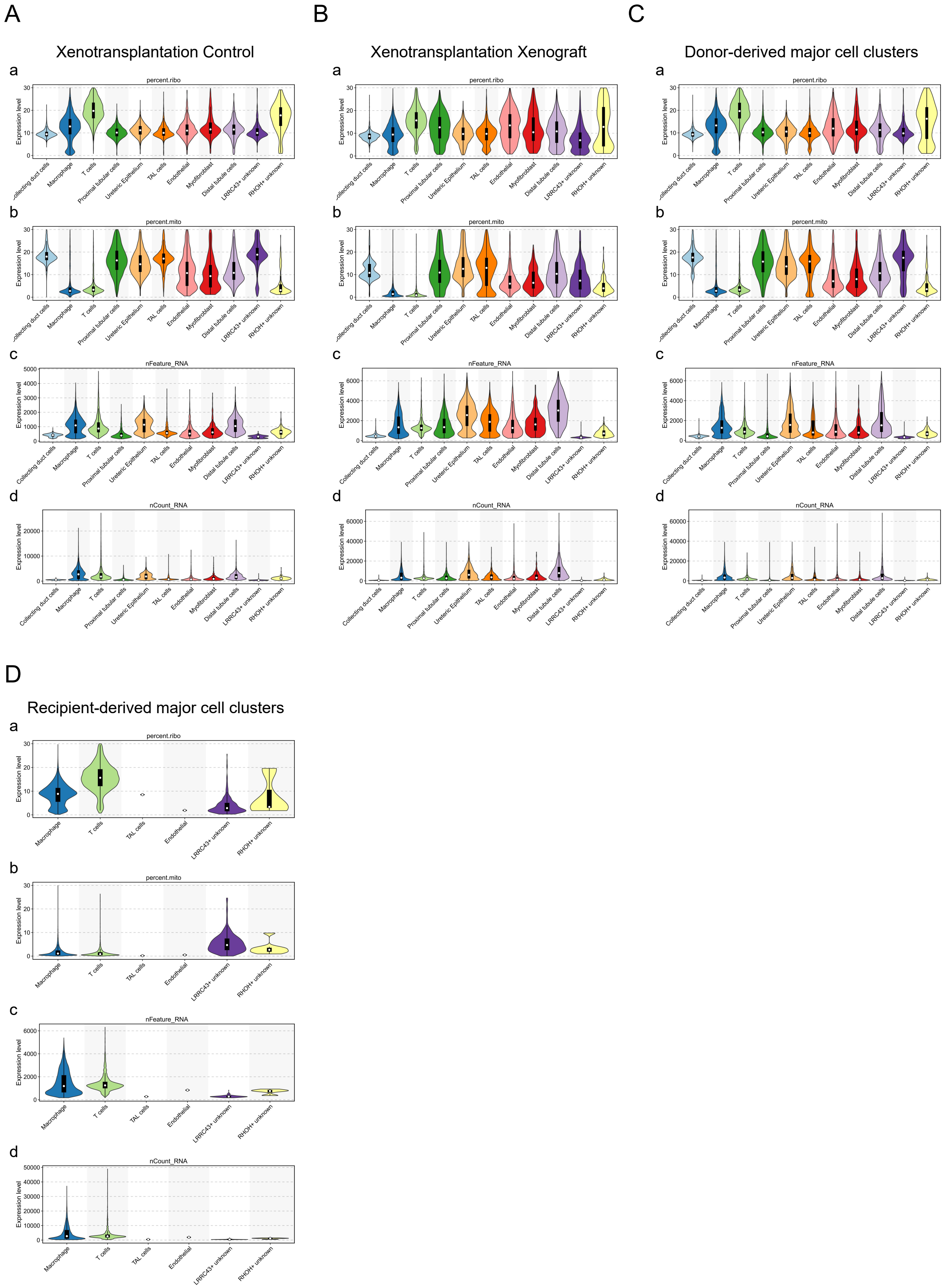

A
Xenotransplantation Control
B
Xenotransplantation Xenograft
C
Donor-derived major cell clusters
D
Recipient-derived major cell clusters
a
percent.ribo
b
percent.mito
c
nFeature_RNA
d
nCount_RNA
Expression level
Collecting duct cells
Macrophage
T cells
Proximal tubular cells
Ureteric Epithelium
TAL cells
Endothelial
Myofibroblast
Distal tubule cells
LRRC43+ unknown
RHOH+ unknown

**Extended Data Figure 1. Violin plots of quality control features for major cell populations across all samples**

(A-B) Quality feature plots of scRNA-seq data of xenotransplantation control and xenograft groups respectively. Each panel including the percentage of ribosomal (ribo) RNA (a), percentage of mitochondrial (mito) RNA (b), total number of expressed genes (nFeature_RNA) (c) and total number of transcripts (nCount_RNA) (d) of each cell.

(C-D) Quality feature plots of scRNA-seq data of donor-derived and recipient-derived major cell clusters respectively. Each panel including the percentage of ribosomal (ribo) RNA (a), percentage of mitochondrial (mito) RNA (b), total number of expressed genes (nFeature_RNA) (c) and total number of transcripts (nCount_RNA) (d) of each cell.

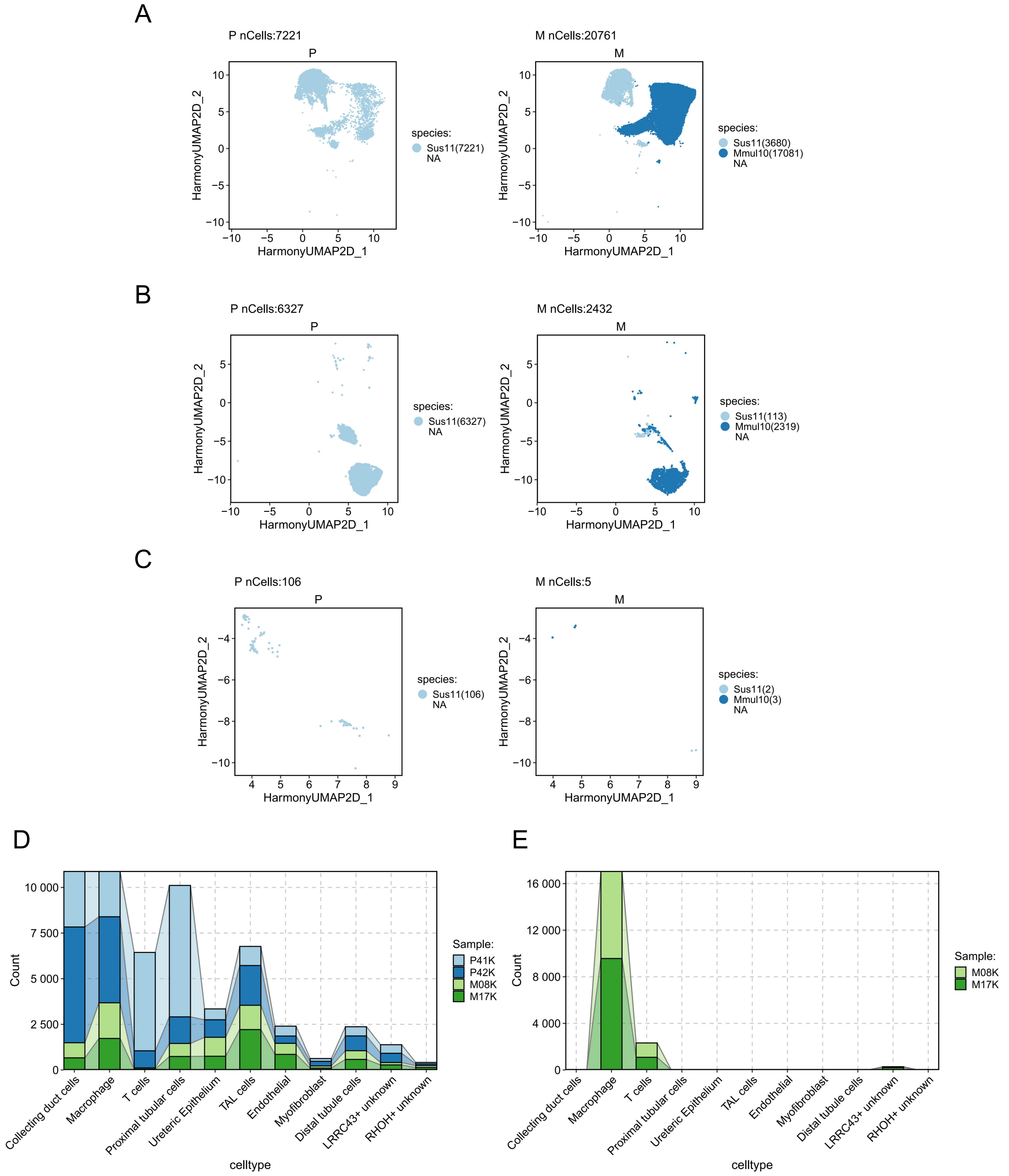
A
P nCells:7221
M nCells:20761
species:
Sus11(7221)
NA
Sus11(3680)
Mmul10(17081)
HarmonyUMAP2D_1
HarmonyUMAP2D_2
B
P nCells:6327
M nCells:2432
Sus11(6327)
Sus11(113)
Mmul10(2319)
C
P nCells:106
M nCells:5
Sus11(106)
Sus11(2)
Mmul10(3)
D
E
Count
celltype
Sample:
P41K
P42K
M08K
M17K
Collecting duct cells
Macrophage
T cells
Proximal tubular cells
Ureteric Epithelium
TAL cells
Endothelial
Myofibroblast
Distal tubule cells
LRRC43+ unknown
RHOH+ unknown

**Extended Data Figure 2. Species assignment and quantification of major immune cell populations in xenografts**

(A) UMAP plot depicting the distribution of cells from different species in marcophage cluster.

(B) UMAP plot depicting the distribution of cells from different species in T cell cluster.

(C) UMAP plot depicting the distribution of cells from different species in B cell cluster.

(D) Quantification of donor-derived major cell counts per individual sample.

(E) Quantification of recipient-derived major cell counts per individual sample.

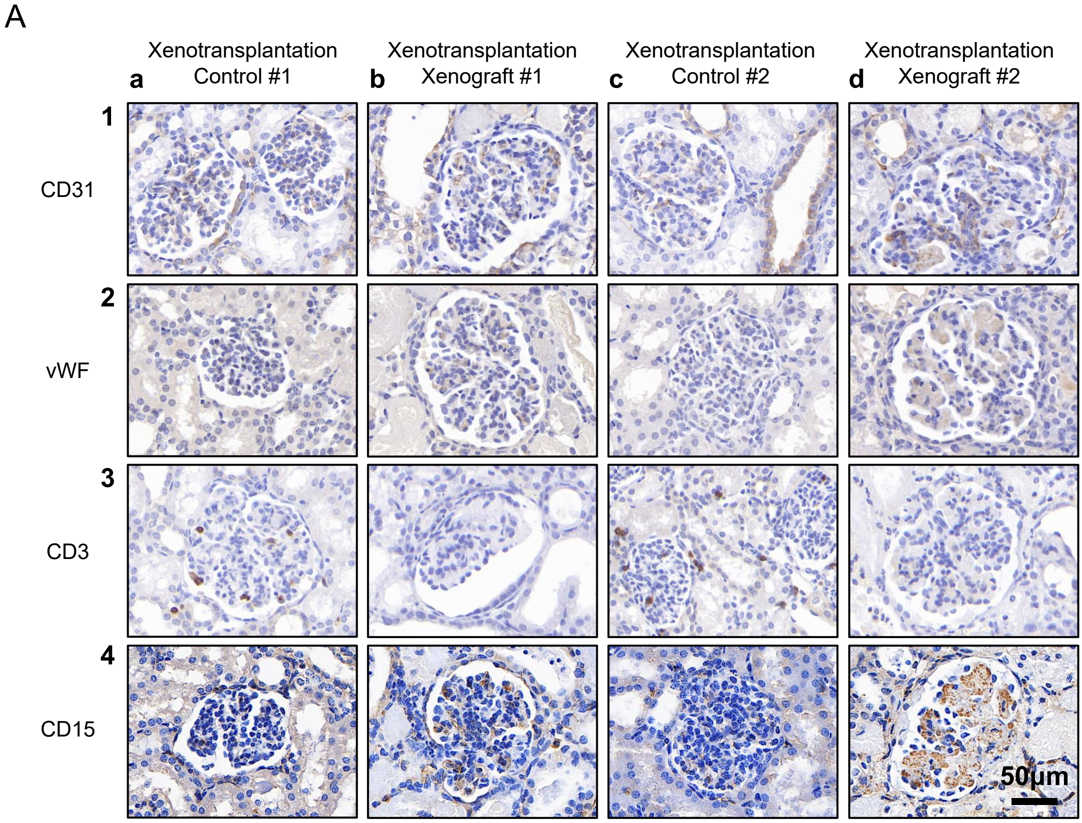
A
Xenotransplantation Control #1
Xenotransplantation Xenograft #1
Xenotransplantation Control #2
Xenotransplantation Xenograft #2
a
b
c
d
1
CD31
2
vWF
3
CD3
4
CD15
50μm

**Extended Data Figure 3. Immunological changes occurring in xenografts after transplantation**

(A) Immunohistochemistry-based evaluation for xenotransplantation control #1 (sections a1-a4) and #2 (sections c1-c4), xenograft #1 (sections b1-b4) and #2 (d1-d4). Fields representative of CD31 (rows 1), vWF (rows 2), CD3 (row 3) and CD15 (row 4) stainings are shown.

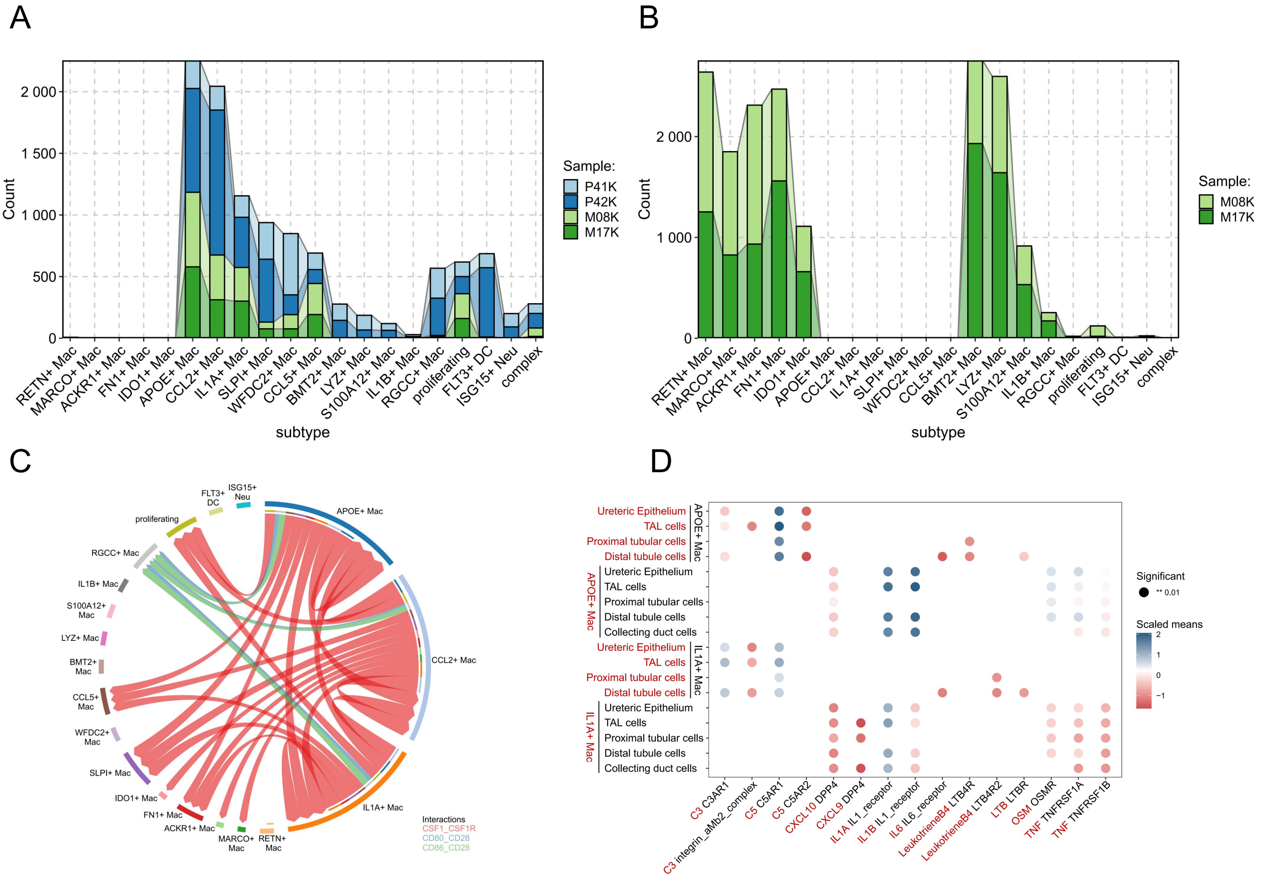
A
B
C
D
Count
subtype
Sample:
P41K
P42K
M08K
M17K
0
500
1 000
1 500
2 000
RETN+ Mac
MARCO+ Mac
ACKR1+ Mac
FN1+ Mac
IDO1+ Mac
APOE+ Mac
CCL2+ Mac
IL1A+ Mac
SLPI+ Mac
WFDC2+ Mac
CCL5+ Mac
BMT2+ Mac
LYZ+ Mac
S100A12+ Mac
IL1B+ Mac
RGCC+ Mac
proliferating
FLT3+ DC
ISG15+ Neu
complex
Interactions
CSF1_CSF1R
CD80_CD28
CD86_CD28
Ureteric Epithelium
TAL cells
Proximal tubular cells
Distal tubule cells
Collecting duct cells
APOE+ Mac
IL1A+ Mac
Significant
** 0.01
Scaled means
C3 C3AR1
C3 integrin_aMb2_complex
C5 C5AR1
C5 C5AR2
CXCL10 DPP4
CXCL9 DPP4
IL1A IL1_receptor
IL1B IL1_receptor
IL6 IL6_receptor
LeukotrieneB4 LTB4R
LeukotrieneB4 LTB4R2
LTB LTBR
OSM OSMR
TNF TNFRSF1A
TNF TNFRSF1B

**Extended Data Figure 4. Myeloid subcluster composition and macrophage-TEC ligand-receptor interactions**

(A) Quantification of donor-derived myeloid cell subclusters counts per individual sample.

(B) Quantification of recipient-derived lmyeloid cell subclusters counts per individual sample.

(C) Circo-plot showing the intercellular interactions between myeloid cell lineages and CCL2+, APOE+ and IL1A+ macrophage subsets. The strings represented interactions determined based on the expression of a ligand by one cell cluster and the expression of a corresponding receptor by another cell cluster. The string thickness reflects the number of different interaction pairs, colored according to cell cluster.

(D) Dot plots showing ligand–receptor interactions between $APOE^{+}$ and $IL1A^{+}$ macrophages and TECs. P values were calculated using an empirical shuffling test.

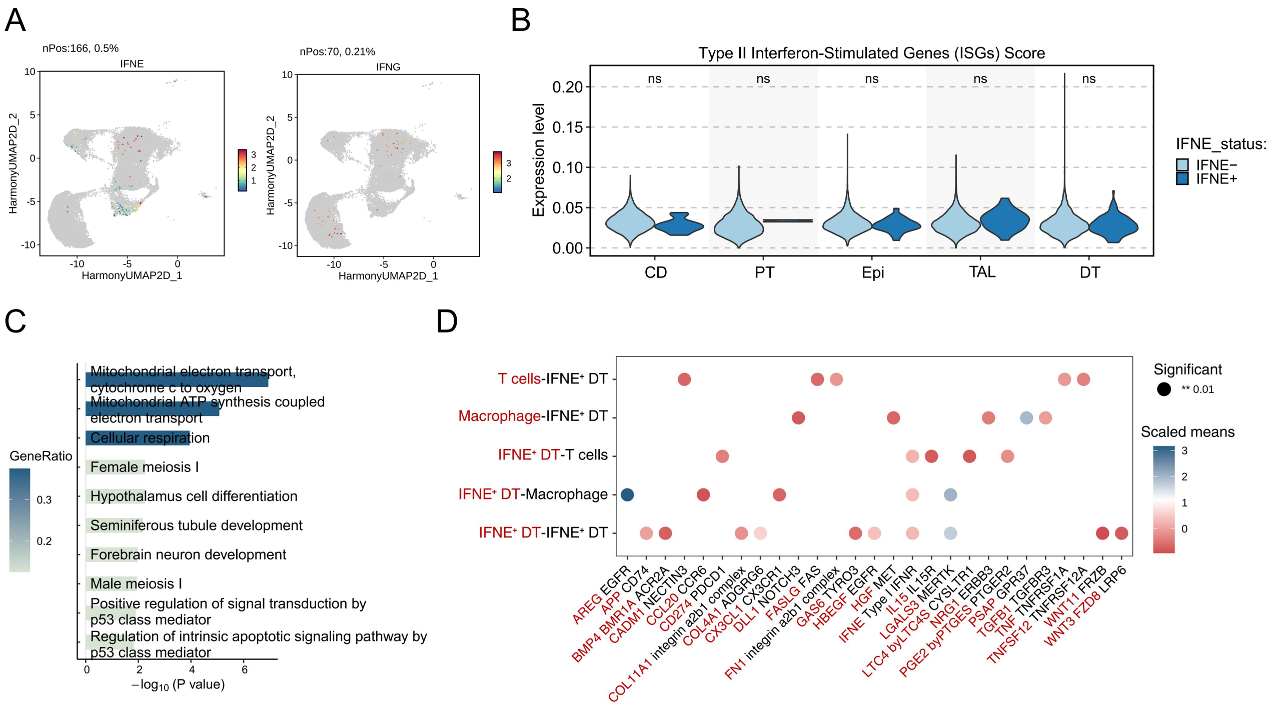
A
nPos:166, 0.5%
IFNE
HarmonyUMAP2D_2
HarmonyUMAP2D_1
nPos:70, 0.21%
IFNG
HarmonyUMAP2D_2
HarmonyUMAP2D_1
B
Type II Interferon-Stimulated Genes (ISGs) Score
Expression level
ns
CD
PT
Epi
TAL
DT
IFNE_status:
IFNE−
IFNE+
C
GeneRatio
Mitochondrial electron transport, cytochrome c to oxygen
Mitochondrial ATP synthesis coupled electron transport
Cellular respiration
Female meiosis I
Hypothalamus cell differentiation
Seminiferous tubule development
Forebrain neuron development
Male meiosis I
Positive regulation of signal transduction by p53 class mediator
Regulation of intrinsic apoptotic signaling pathway by p53 class mediator
−log10 (P value)
D
T cells-IFNE+ DT
Macrophage-IFNE+ DT
IFNE+ DT-T cells
IFNE+ DT-Macrophage
IFNE+ DT-IFNE+ DT
Significant
** 0.01
Scaled means
AREG EGFR
APP CD74
BMP4 BMR1A ACR2A
CADM1 NECTIN3
CCL20 CCR6
CD274 PDCD1
COL11A1 integrin a2b1 complex
COL4A1 ADGRG6
CX3CL1 CX3CR1
DLL1 NOTCH3
FASLG FAS
FN1 integrin a2b1 complex
GAS6 TYRO3
HBEGF EGFR
HGF MET
IFNE Type I IFNR
IL15 IL15R
LGALS3 MERTK
LTC4 byLTC4S CYSLTR1
NRG1 ERBB3
PGE2 byPTGES PTGER2
PSAP GPR37
TGFB1 TGFBR3
TNF TNFRSF1A
TNFSF12 TNFRSF12A
WNT11 FRZB
WNT3 FZD8 LRP6

**Extended Data Figure 5. IFNE⁺ tubular cells engage in immune crosstalk**
(A) UMAP visualization of IFNE and IFNG expression across major cell clusters.
(B) Type II ISG module scores in IFNE+ versus IFNE- TECs across all clusters.
(C) GO enrichment analysis of genes downregulated in IFNE+ TECs compared to IFNE- TECs.
(D) Dot plots depicting ligand-receptor interactions between $IFNE^{+}$ distal tubule cells and T cells, as well as between macrophages and $IFNE^{+}$ distal tubule cells. P values were calculated using an empirical shuffling test.

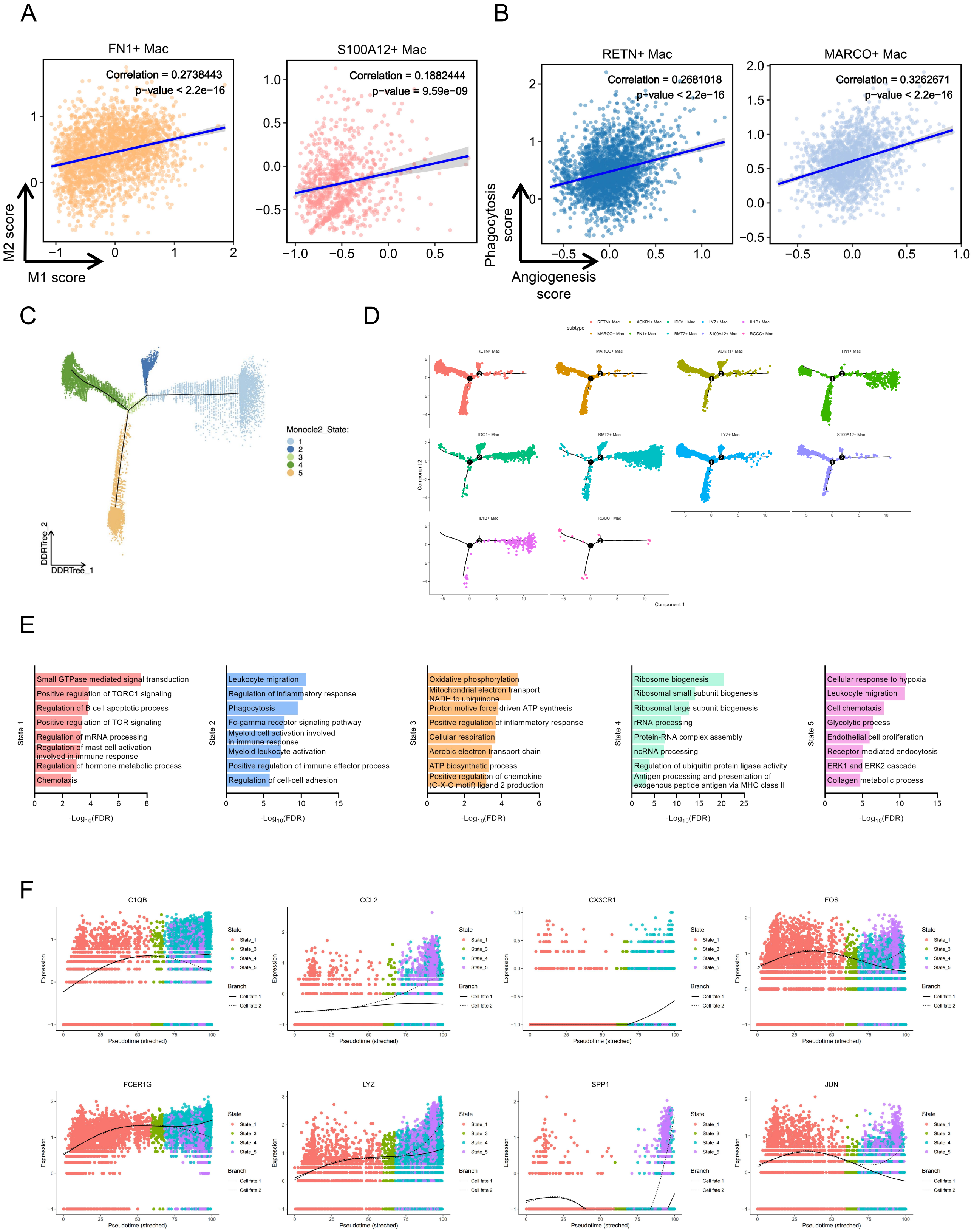
A
FN1+ Mac
Correlation = 0.2738443
p−value < 2.2e−16
M2 score
M1 score
S100A12+ Mac
Correlation = 0.1882444
p−value = 9.59e−09
B
RETN+ Mac
Correlation = 0.2681018
p−value < 2.2e−16
Phagocytosis score
Angiogenesis score
MARCO+ Mac
Correlation = 0.3262671
p−value < 2.2e−16
C
Monocle2_State:
1
2
3
4
5
DDRTree_2
DDRTree_1
D
subtype
RETN+ Mac
ACKR1+ Mac
IDO1+ Mac
LYZ+ Mac
IL1B+ Mac
MARCO+ Mac
FN1+ Mac
BMT2+ Mac
S100A12+ Mac
RGCC+ Mac
Component 2
Component 1
E
State 1
Small GTPase mediated signal transduction
Positive regulation of TORC1 signaling
Regulation of B cell apoptotic process
Positive regulation of TOR signaling
Regulation of mRNA processing
Regulation of mast cell activation involved in immune response
Regulation of hormone metabolic process
Chemotaxis
State 2
Leukocyte migration
Regulation of inflammatory response
Phagocytosis
Fc-gamma receptor signaling pathway
Myeloid cell activation involved in immune response
Myeloid leukocyte activation
Positive regulation of immune effector process
Regulation of cell-cell adhesion
State 3
Oxidative phosphorylation
Mitochondrial electron transport NADH to ubiquinone
Proton motive force-driven ATP synthesis
Positive regulation of inflammatory response
Cellular respiration
Aerobic electron transport chain
ATP biosynthetic process
Positive regulation of chemokine (C-X-C motif) ligand 2 production
State 4
Ribosome biogenesis
Ribosomal small subunit biogenesis
Ribosomal large subunit biogenesis
rRNA processing
Protein-RNA complex assembly
ncRNA processing
Regulation of ubiquitin protein ligase activity
Antigen processing and presentation of exogenous peptide antigen via MHC class II
State 5
Cellular response to hypoxia
Leukocyte migration
Cell chemotaxis
Glycolytic process
Endothelial cell proliferation
Receptor-mediated endocytosis
ERK1 and ERK2 cascade
Collagen metabolic process
-Log10(FDR)
F
C1QB
CCL2
CX3CR1
FOS
FCER1G
LYZ
SPP1
JUN
Expression
Pseudotime (streched)
State
State_1
State_3
State_4
State_5
Branch
Cell fate 1
Cell fate 2

**Extended Data Figure 6. Pseudotemporal dynamics of recipient macrophage polarization and function.**

(A) Scatter plot depicting the Pearson correlation coefficient between M1 and M2 signature scores in $FN1^+$ and $S100A12^+$ recipient-derived macrophage subsets.

(B) Scatter plot depicting the Pearson correlation coefficient between phagocytosis and angiogenesis signature scores in $RETN^+$ and $MARCO^+$ recipient-derived macrophage subsets.

(C) Pseudotime analysis reveals 5 distinct cell state clusters, shown in different colors on the trajectory plot. Each dot represents a single cell, with its color reflecting the cell state category (State 1 to State 5).

(D) Distribution of subsets of recipient-derived macrophages along the pseudotime differentiation trajectory.

(E) GO enrichment analysis of differentially expressed genes in 5 distinct cell state clusters.

(F) Two-dimensional plot showing the dynamic expression of representative genes during the recipient-derived macrophages transitions along the pseudotime.

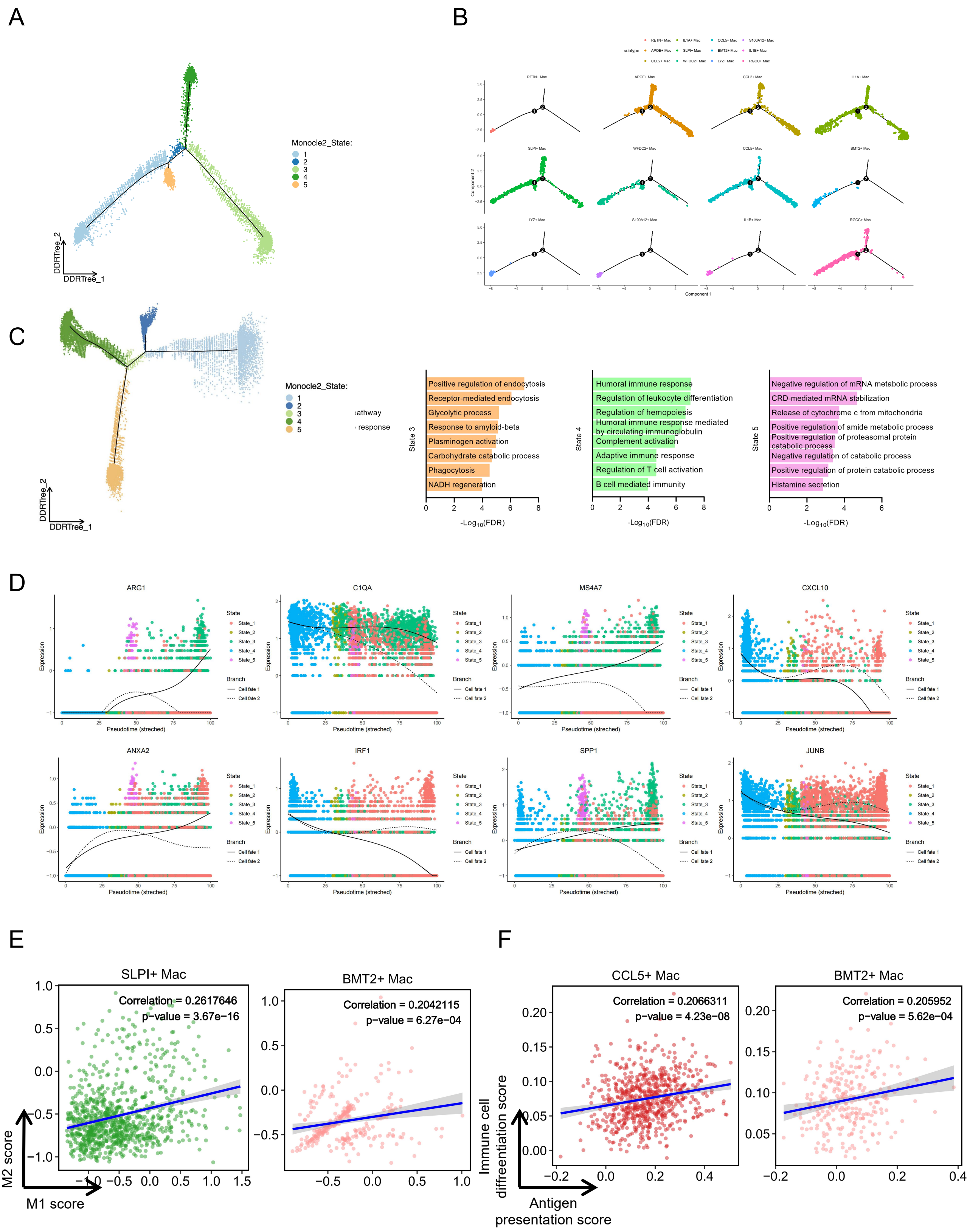
A
B
C
D
E
F
Monocle2_State:
DDRTree_1
DDRTree_2
Component 1
Component 2
State 3
Positive regulation of endocytosis
Receptor-mediated endocytosis
Glycolytic process
Response to amyloid-beta
Plasminogen activation
Carbohydrate catabolic process
Phagocytosis
NADH regeneration
State 4
Humoral immune response
Regulation of leukocyte differentiation
Regulation of hemopoiesis
Humoral immune response mediated by circulating immunoglobulin
Complement activation
Adaptive immune response
Regulation of T cell activation
B cell mediated immunity
State 5
Negative regulation of mRNA metabolic process
CRD-mediated mRNA stabilization
Release of cytochrome c from mitochondria
Positive regulation of amide metabolic process
Positive regulation of proteasomal protein catabolic process
Negative regulation of catabolic process
Positive regulation of protein catabolic process
Histamine secretion
$-Log_{10}(FDR)$
ARG1
C1QA
MS4A7
CXCL10
ANXA2
IRF1
SPP1
JUNB
Expression
Pseudotime (streched)
State
Branch
Cell fate 1
Cell fate 2
SLPI+ Mac
Correlation = 0.2617646
p−value = 3.67e−16
BMT2+ Mac
Correlation = 0.2042115
p−value = 6.27e−04
M2 score
M1 score
CCL5+ Mac
Correlation = 0.2066311
p−value = 4.23e−08
BMT2+ Mac
Correlation = 0.205952
p−value = 5.62e−04
Immune cell differentiation score
Antigen presentation score

**Extended Data Figure 7. Resident macrophages exhibit divergent trajectories and functional plasticity.**

(A) Pseudotime analysis reveals 5 distinct cell state clusters, shown in different colors on the trajectory plot. Each dot represents a single cell, with its color reflecting the cell state category (State 1 to State 5).

(B) Distribution of subsets of tissue-resident macrophages along the pseudotime differentiation trajectory.

(C) GO enrichment analysis of differentially expressed genes in 5 distinct cell state clusters.

(D) Two-dimensional plot showing the dynamic expression of representative genes during the tissue-resident macrophages transitions along the pseudotime.

(E) Scatter plot depicting the Pearson correlation coefficient between M1 and M2 signature scores in $SLPI^+$ and $BMT2^+$ tissue-resident macrophage subsets.

(F) Scatter plot depicting the Pearson correlation coefficient between phagocytosis and angiogenesis signature scores in $CCL5^+$ and $BMT2^+$ tissue-resident macrophage subsets.

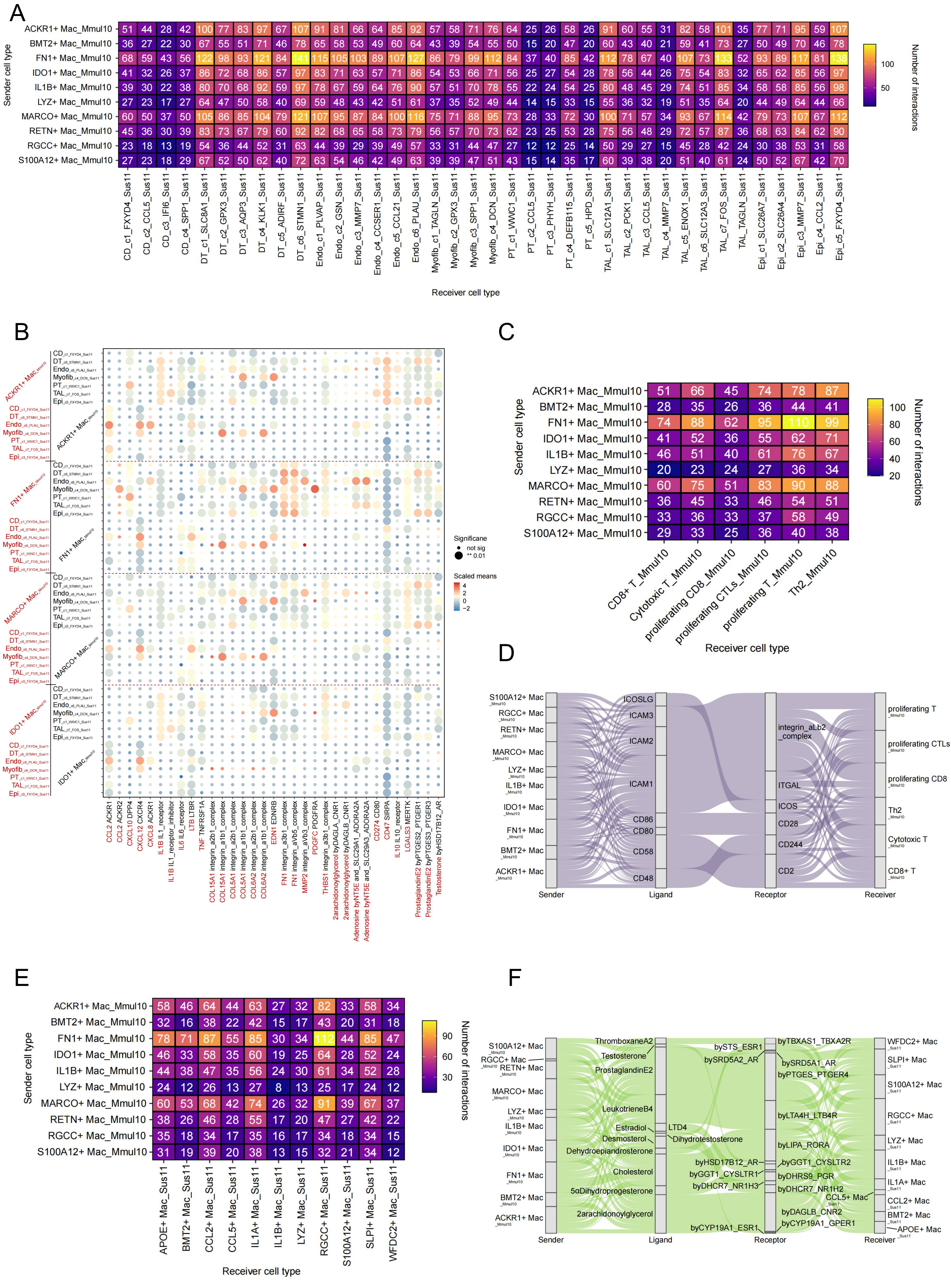
A
Sender cell type
ACKR1+ Mac_Mmul10
BMT2+ Mac_Mmul10
FN1+ Mac_Mmul10
IDO1+ Mac_Mmul10
IL1B+ Mac_Mmul10
LYZ+ Mac_Mmul10
MARCO+ Mac_Mmul10
RETN+ Mac_Mmul10
RGCC+ Mac_Mmul10
S100A12+ Mac_Mmul10
Number of interactions
Receiver cell type
B
Significance
not sig
** 0.01
Scaled means
C
Sender cell type
CD8+ T_Mmul10
Cytotoxic T_Mmul10
proliferating CD8_Mmul10
proliferating CTLs_Mmul10
proliferating T_Mmul10
Th2_Mmul10
Receiver cell type
Number of interactions
D
Sender
Ligand
Receptor
Receiver
E
Sender cell type
APOE+ Mac_Sus11
BMT2+ Mac_Sus11
CCL2+ Mac_Sus11
CCL5+ Mac_Sus11
IL1A+ Mac_Sus11
IL1B+ Mac_Sus11
LYZ+ Mac_Sus11
RGCC+ Mac_Sus11
S100A12+ Mac_Sus11
SLPI+ Mac_Sus11
WFDC2+ Mac_Sus11
Receiver cell type
Number of interactions
F
Sender
Ligand
Receptor
Receiver

**Extended Data Figure 8. Intercellular interactions between macrophages and kidney cells.**

(A) Heatmap showing numbers of potential ligand–receptor (L-R) pairs between recipient-derived macrophage subsets and kidney cell subsets.
(B) Dot plots showing L-R interactions between selected recipient-derived macrophage subsets and kidney cell subsets. P values were calculated using an empirical shuffling test.
(C) Heatmap showing numbers of potential L-R pairs between recipient-derived macrophage subsets and T cell subsets.
(D) Sankey diagram showing the pro-T cell activating-specific interactions recipient-derived macrophage subsets and T cell subsets.
(E) Heatmap showing numbers of potential L-R pairs between recipient-derived macrophage subsets and tissue-resident macrophage subsets.
(F) Sankey diagram showing the lipid molecules-specific interactions between recipient-derived macrophage subsets and tissue-resident macrophage subsets.

**Extended Data Figure 9. T cell dynamics and immune-stromal interactions.**
(A) Quantification of donor-derived lymphoid cell subclusters counts per individual sample.
(B) Quantification of recipient-derived lymphoid cell subclusters counts per individual sample.
(C) Pseudotime analysis reveals 7 distinct cell state clusters, shown in different colors on the trajectory plot. Each dot represents a single cell, with its color reflecting the cell state category (State 1 to State 7).
(D) Distribution of subsets of recipient-derived T cells along the pseudotime differentiation trajectory.
(E) GO enrichment analysis of differentially expressed genes in 7 distinct cell state clusters.
(F) Heatmap depicting the dynamic expression changes of genes and related pathways of recipient-derived T cells with different cell fate along pseudotime.
(G) Two-dimensional plot showing the dynamic expression of representative genes during the recipient-derived T cells transitions along the pseudotime.
(H) Heatmap showing numbers of potential ligand–receptor pairs between recipient-derived T cells subsets and kidney cell subsets.
(I) Sankey diagram showing the WNT8B interactors between recipient-derived proliferating T cells and selected kidney cells subsets.
(J) Sankey diagram showing the CD40LG interactors between recipient-derived Th2 cells and selected kidney cells subsets.
